# Nanopore ReCappable Sequencing maps SARS-CoV-2 5′ capping sites and provides new insights into the structure of sgRNAs

**DOI:** 10.1101/2021.11.24.469860

**Authors:** Camilla Ugolini, Logan Mulroney, Adrien Leger, Matteo Castelli, Elena Criscuolo, Maia Kavanagh Williamson, Andrew D Davidson, Abdulaziz Almuqrin, Roberto Giambruno, Miten Jain, Gianmaria Frige, Hugh Olsen, George Tzertzinis, Ira Schildkraut, Madalee G. Wulf, Ivan R. Corrêa, Laurence Ettwiller, Nicola Clementi, Massimo Clementi, Nicasio Mancini, Ewan Birney, Mark Akeson, Francesco Nicassio, David A. Matthews, Tommaso Leonardi

**Affiliations:** Center for Genomic Science of IIT@SEMM, Fondazione Istituto Italiano di Tecnologia, 20139 Milano, Italy; European Molecular Biology Laboratory, European Bioinformatics Institute, Hinxton, UK; Biomolecular Engineering Department, UC Santa Cruz, CA 95064, USA; Laboratory of Microbiology and Virology, Vita-Salute San Raffaele University; via Olgettina 58, 20132 Milan, Italy; School of Cellular and Molecular Medicine, Faculty of Life Sciences, University Walk, University of Bristol, Bristol, BS8 1TD, UK; Department of Clinical Laboratory Sciences, King Saud University, Riyadh, Saudi Arabia; Department of Experimental Oncology, European Institute of Oncology IRCCS, 20132 Milano, Italy; New England Biolabs, Ipswich, MA 01938 3, USA; Laboratory of Medical Microbiology and Virology, IRCCS San Raffaele Scientific Institute; via Olgettina 60, 20132 Milan, Italy

## Abstract

The SARS-CoV-2 virus has a complex transcriptome characterised by multiple, nested sub genomic RNAs used to express structural and accessory proteins. Long-read sequencing technologies such as nanopore direct RNA sequencing can recover full-length transcripts, greatly simplifying the assembly of structurally complex RNAs. However, these techniques do not detect the 5′ cap, thus preventing reliable identification and quantification of full-length, coding transcript models. Here we used Nanopore ReCappable Sequencing (NRCeq), a new technique that can identify capped full-length RNAs, to assemble a complete annotation of SARS-CoV-2 sgRNAs and annotate the location of capping sites across the viral genome. We obtained robust estimates of sgRNA expression across cell lines and viral isolates and identified novel canonical and non-canonical sgRNAs, including one that uses a previously un-annotated leader-to-body junction site. The data generated in this work constitute a useful resource for the scientific community and provide important insights into the mechanisms that regulate the transcription of SARS-CoV-2 sgRNAs.

## Introduction

SARS-CoV-2 is an enveloped virus with a ∼30kb long, positive-sense single-stranded RNA genome^1^ that belongs to the family of *Coronaviridae* in the *Nidovirales* order. All family members share the same genomic architecture, that consists of a capped 5′ untranslated region (UTR) containing a leader transcription regulatory sequence (TRS-L), a large open reading frame (ORF) – ORF1ab – encoding for a single polyprotein that self-cleaves into the non-structural proteins (Nsps), followed by multiple ORFs encoding for the structural and accessory proteins and a polyadenylated 3′ UTR. Except for 1ab, each ORF is preceded by a body TRS (TRS-B) highly homologous to the TRS-L. These genomic features are at the basis of coronaviruses (CoVs) regulation of protein abundance and timely expression. Nsps are directly translated from CoVs genome in the early phases of the replicative cycle to assemble the replication-transcription complex (RTC), while all other ORFs are translated from subgenomic RNAs (sgRNAs) that are subsequently synthesized by the RTC through negative sense intermediates.

According to the prevailing model, during the synthesis of the negative stranded intermediates the RTC can undergo a template switching event at each TRS-B sequence encountered^2–5^. This leads to the production of intermediates of different lengths formed by the fusion of the 5′-UTR TRS-L sequence with a TRS-B immediately upstream of the various ORFs encoded by the viral genome. The result of this process is a landscape of negative-strand, partially overlapping sgRNAs with length ranging from ∼200nt to over 8000nt (**Figure 1A**). These sgRNA intermediates are then used as a template for the synthesis of positive-strand, protein coding sgRNAs, that are 5′-capped by the viral machinery and in turn translated to produce structural and accessory proteins^4,6,7^. This structural complexity of the SARS-CoV-2 transcriptome poses real challenges for transcriptome assembly and sgRNA quantification, particularly for short read sequencing technologies. To overcome these limitations, recent works have used long-read sequencing techniques such as PacBio SMRT and Nanopore direct RNA sequencing (DRS) to reconstruct the transcriptional architecture of SARS-CoV-2^4,6–10^. DRS is a technique that measures ionic current alterations produced by the translocation of nucleotides through a nanopore, which theoretically allows RNA molecules of any length to be sequenced as they exist in the cell, without the need for retrotranscription or amplification^11,12^. Despite the advantages of DRS over short-read cDNA sequencing, this technique is still unable to differentiate full-length transcripts from RNA fragments resulting from RNA degradation or incomplete sequencing^11^. This limitation poses a real challenge for studying SARS-CoV-2 sgRNAs, because it makes it impossible to discriminate between *bona fide*, capped, non-canonical sgRNAs that lack a leader-to-body fusion from uncapped RNA fragments or incompletely sequenced reads. Furthermore, RNA degradation leads to a large number of reads mapping to the 3′ region common to all sgRNAs, thus introducing a significant bias when trying to quantify sgRNA expression.

**Figure 1.**
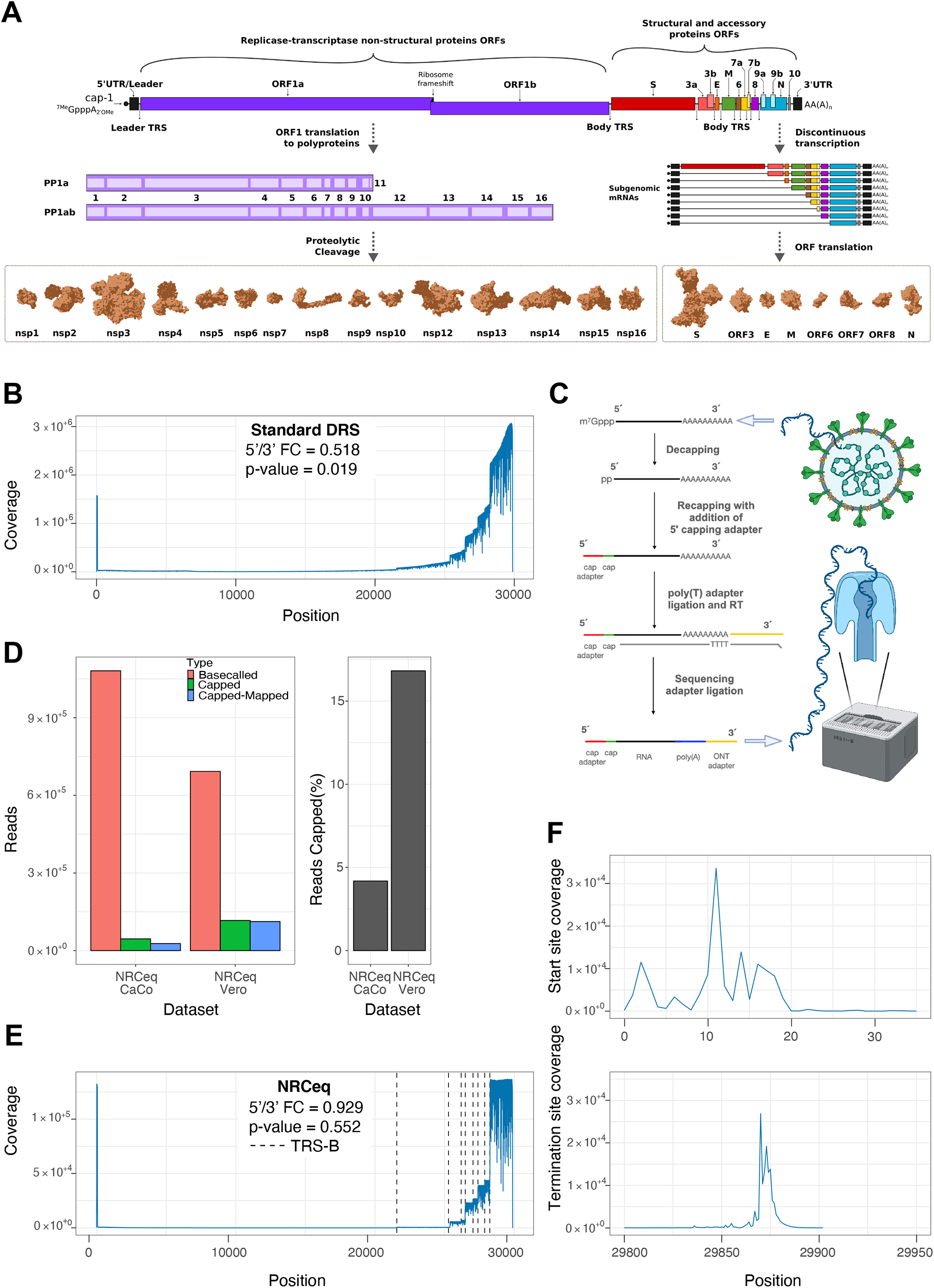
NRCeq allows sequencing of full-length viral sgRNAs. (**A**) Schematic representation of the landscape of the SARS-CoV-2 transcriptome, ORFs cleavage sites and protein structures^42^. (**B**) Read coverage across the viral genome calculated from the aggregated standard Nanopore DRS datasets used in this study (see Suppl. Table 1). The figure also reports the coverage fold change between the 5′ (from 15 to 60) and 3′ (region from 29805 to 29850). The reported p-value was calculated with the two-sided Welch’s test. (**C**) Schematic overview of the NRCeq recapping protocol. (**D**) Number (left) and percentage (right) of basecalled, trimmed and mapped reads for the NRCeq datasets. (**E**) Read coverage across the viral genome calculated using aggregated NRCeq data from CaCo2 and Vero cells.The figure also reports the coverage fold change between the 5′ (from 15 to 60) and 3′ (region from 29805 to 29850). The reported p-value was calculated with the two-sided Welch’s test. (**F**) Coverage of the viral genome calculated using only the alignment start sites (top) or alignment termination sites (bottom) for the NRCeq data from CaCo2 and Vero cells, aggregated in a single dataset.

The recent development of Nanopore ReCappable Sequencing (NRCeq) overcomes this limitation by replacing the native 5′ cap with a 5′ cap-linked RNA sequencing adapter, allowing to discriminate full-length, capped molecules from fragmented RNAs and truncated sequencing artefacts^13^. Using this approach we assembled a de novo SARS-CoV-2 transcriptome, that includes well-supported canonical and non-canonical transcripts that encode the ORFs annotated in Uniprot^14^. The assembly has been refined by bioinformatic pipelines which gave insights into the presence of deletions in sgRNAs, genuine capped non-canonical sgRNAs as well as a novel ORF.

We also show that quantifying standard DRS datasets against the NRCeq transcriptome assembly provides robust expression estimates of sgRNAs across multiple cell lines and viral strains.

## Results

### NRCeq recovers full-length reads

In order to assemble a comprehensive SARS-CoV-2 transcriptome we generated six Nanopore direct RNA sequencing (DRS) datasets from Vero, CaCo2 and CaLu3 cells infected with SARS-CoV-2 and analyzed them in combination with publicly available DRS data (see **Suppl. Table 1** and **Materials and Methods**). We processed the DRS datasets with a custom pipeline (**Suppl. Fig. 1**) that performed quality control and mapping to the reference viral genome (see **Materials and Methods**). In line with current models of SARS-CoV-2 discontinuous transcription, we expected similar levels of coverage at the 5′ region (upstream of the TRS-L) and the 3′ region (downstream of the last TRS-B), because these two regions are common to all known sgRNAs. However, we observed that the coverage was significantly higher at the 3′ end (5′/3′ coverage Fold Change 0.518, p-value=0.019, **Fig. 1B**) and that it gradually decreased from 3′ to 5′. Because DRS starts from the polyA tail and proceeds in the 3′ to 5′ direction, we reasoned that such a discrepancy in coverage could be explained by sgRNAs that lacked the leader sequence or incomplete sequencing reads and RNA degradation fragments^11,13,15,16^. Both explanations would likely result in ambiguous assignment of the incomplete reads to multiple sgRNAs confounding expression estimates. To overcome this limitation we used NRCeq, a recent DRS protocol that specifically couples an RNA adapter to m^7^G capped 5′ ends, which permits identification of *bona fide* full-length RNA reads^13^. We infected CaCo2 and Vero cells with SARS-CoV-2 (strain in **Suppl. Table 1**) and generated NRCeq libraries for nanopore DRS (**Fig. 1C** and see **Methods**). On average, we achieved a recapping efficiency of 10.5% (**Fig. 1D**), with an average of 78.2% of cap-adapted reads mapping to the viral genome. We observed a difference in the number of viral reads between CaCo2 and Vero cells (respectively 60.1% and 96.3% of mapped reads over cap adapted reads, **Fig. 1D**), which is in agreement with previous reports showing different viral titers in the two cell lines^17^.

After aligning the NRCeq datasets to the SARS-CoV-2 reference genome, we observed similar levels of coverage for the region upstream of the TRS-L and at the 3′ end of the viral genome (FC=0.929, p-value=0.552 **Fig. 1E**). Additionally, there was uniform coverage across the body of each annotated ORF, with sharp drops in coverage near each TRS-B sequence reflecting the RTC template switching events. We found that 95.7% of the alignments start at genomic positions between 0-30 and 94.5% of the alignments end at genomic positions between 29860-29890 (**Fig. 1F and Suppl. Fig. 2**). These observations support the accepted model of SARS-CoV-2 discontinuous transcription, where most of the viral sgRNAs are the result of the fusion of an ORF with the 5′ leader sequence.

### Genome wide identification of 5′ capping sites

We then used the NRCeq data to obtain a genome-wide map of the 5′ capped starting sites of sgRNAs. To this end, we performed a peak calling analysis using only the 5′ nucleotide of the alignment of each NRCeq mapped read (**Suppl. Table 2**, see **Materials and Methods**). We identified two major peaks at genomic coordinates 6 and 13, plus two smaller peaks (at 36 and 44) that represent alignments with an incomplete 5′ UTR. The peaks at 6 and 13 likely correspond to sgRNAs with a full length 5′ UTR that was incompletely aligned due to the miscall of the molecule terminal nucleotides typical of DRS^11–13^. Other minor peaks supported by varying numbers of reads (reported in **Suppl. Table 2**) were also observed in the regions between 27300-28300 and at genomic coordinate 9193, representing low abundance non-canonical transcripts or alignments artefacts (**Suppl. Fig. 3** and **Suppl. Table 2**). In contrast, when we performed the 5′ peak calling using standard DRS datasets, we observed a highly fragmented profile along the entire genome, making it impossible to discern real 5′ cap sites from 5′ ends generated by fragmentation events (**Suppl. Fig. 3** and **Suppl. Tracks**). All together, these observations support NRCeq’s ability to capture full-length, m^7^G capped viral sgRNAs, allowing accurate investigation of the complex, nested transcriptional landscape of SARS-CoV-2.

### Assembly of a comprehensive SARS-CoV-2 transcriptome

Thanks to the improved confidence in capturing full-length transcripts, we could use the NRCeq datasets to assemble full-length transcript models for SARS-CoV2 sgRNAs. To this end, we developed a pipeline that basecalls the raw NRCeq data, trims the 5′ NRCeq adapter, maps the reads to the SARS-CoV-2 genome and assembles transcript models (see **Materials and Methods**). Briefly, the pipeline uses Guppy for basecalling, Porechop to identify the cap-adapted reads and trim the 5′ adapter sequence, minimap2^18^ to align the reads to the reference genome, and Pinfish to assemble the transcriptome. This resulted in a consensus assembly of 21 transcript models (**Fig. 2A** and **Suppl. Table 3**), 14 of which we could classify as canonical based on the presence of: 1) a start site aligning to the 5′ end of the genome; 2) a 5′ UTR of at least 40 nucleotides; 3) a termination site aligning to the 3′ of the viral genome; and 4) a body-to-leader fusion in a region with at least 66.0% similarity to the canonical TRS-B sequence (see **Materials and Methods**). The assembled transcript models have length ranging from 370 to 8374nt; 6 of them are derived from a single, contiguous genomic region, 13 from two discontinuous regions and 2 from three. (**Suppl. Fig. 4 A-B-C**).

**Figure 2.**
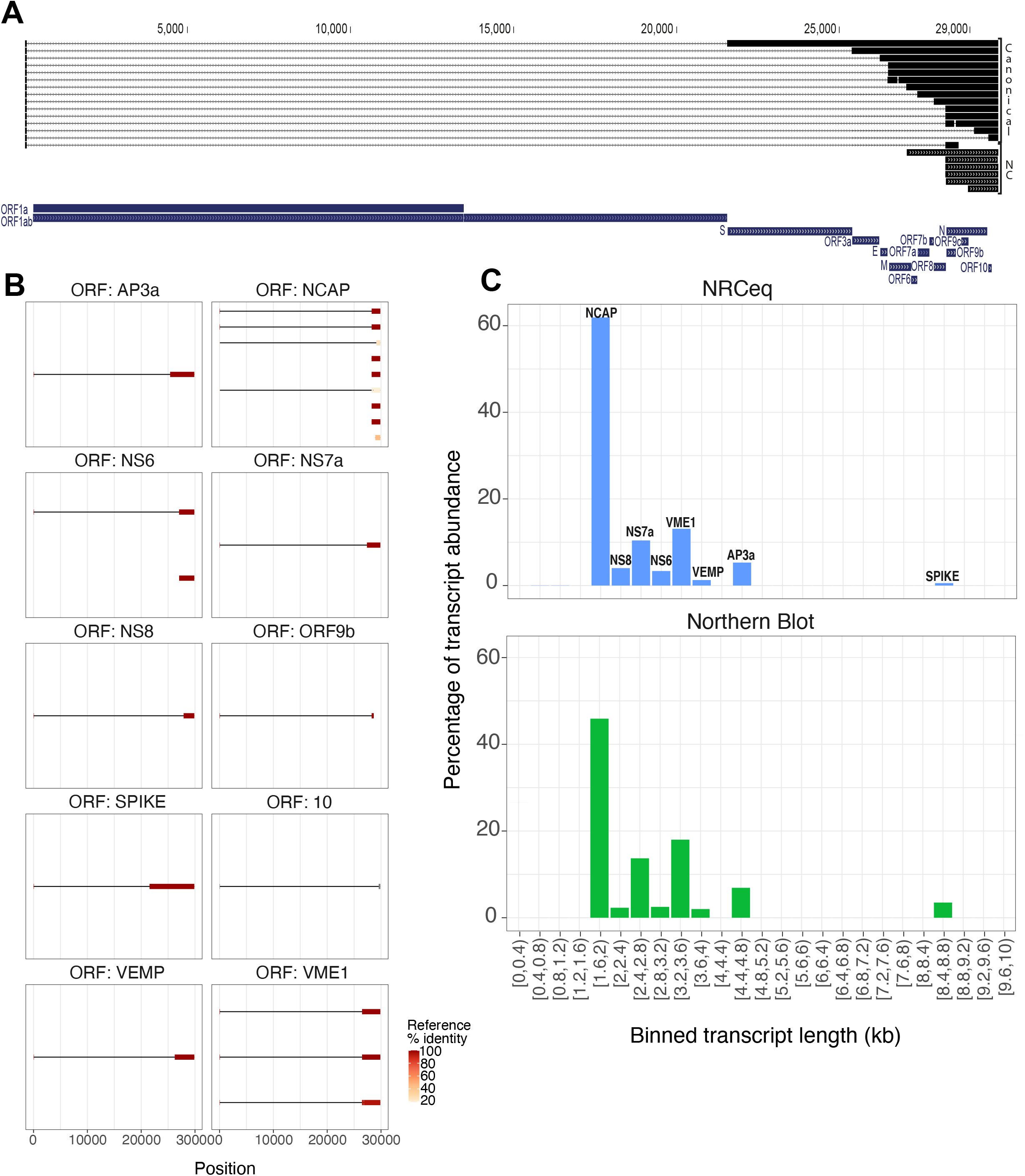
NRCeq assembly identifies and quantifies viral sgRNAs. (**A**) UCSC Genome Browser track showing the SARS-CoV-2 transcriptome assembly obtained with NRCeq data. The figure reports both canonical and non-canonical (NC) transcript models. SARS-CoV-2 ORFs are reported for reference. (**B**) Diagram showing the structure of each assembled transcript, subdivided according to the ORF encoded. The color scale indicates the number of identical amino acids between the first ORF of the sgRNA and the best match in the reference SARS-CoV2 proteome (Uniprot) expressed as a fraction of the reference protein’s length. (**C**) Quantification of the ORFs performed by NRCeq and Northern Blot. NRCeq data from CaCo2 and Vero cells were aggregated in a single dataset. For each bin of 400nt (x-axis) the cumulative expression of all assembled transcript models was calculated and expressed as a percentage (y-axis). The Northern Blot quantification data was obtained from Ogando et al.^22^

We then assigned each transcript model to a specific sgRNA based on the sequence homology between the first encoded ORF and the viral proteome^14^. At least one transcript model was assigned to each annotated ORF (S, 3a, E, M, 6, 7a/b, 8, N, 10, **Materials and Methods, Fig. 2B, Suppl. Fig. 4D**). Notably, our assembly also included sgRNA models for ORF10, the existence of which was the recent subject of discordant reports^4,8,19^, as well a new ORF internal to N, which we named ORF9d. Importantly, NRCeq reads supporting ORF10 and ORF9d were identified both in CaCo2 and Vero cells.

### ORF10 and ORF9d are encoded by canonical, capped sgRNAs

The NRCeq assembly included a canonical transcript model encoding ORF10, which is supported by 116 reads, each with a junction between the canonical TRS-L and one of several non-canonical (i.e. with 1 or 2 mismatches) TRS-B sequences located in the interval ∼29,300-29,700. These reads contain 2 possible start codons in-frame with the annotated ORF10 and both are within ∼15nt of ribosomal footprinting peaks. This supports the existence of functional, translated ORF10 sgRNAs as previously reported^19^ (**Fig 3B**).

**Figure 3.**
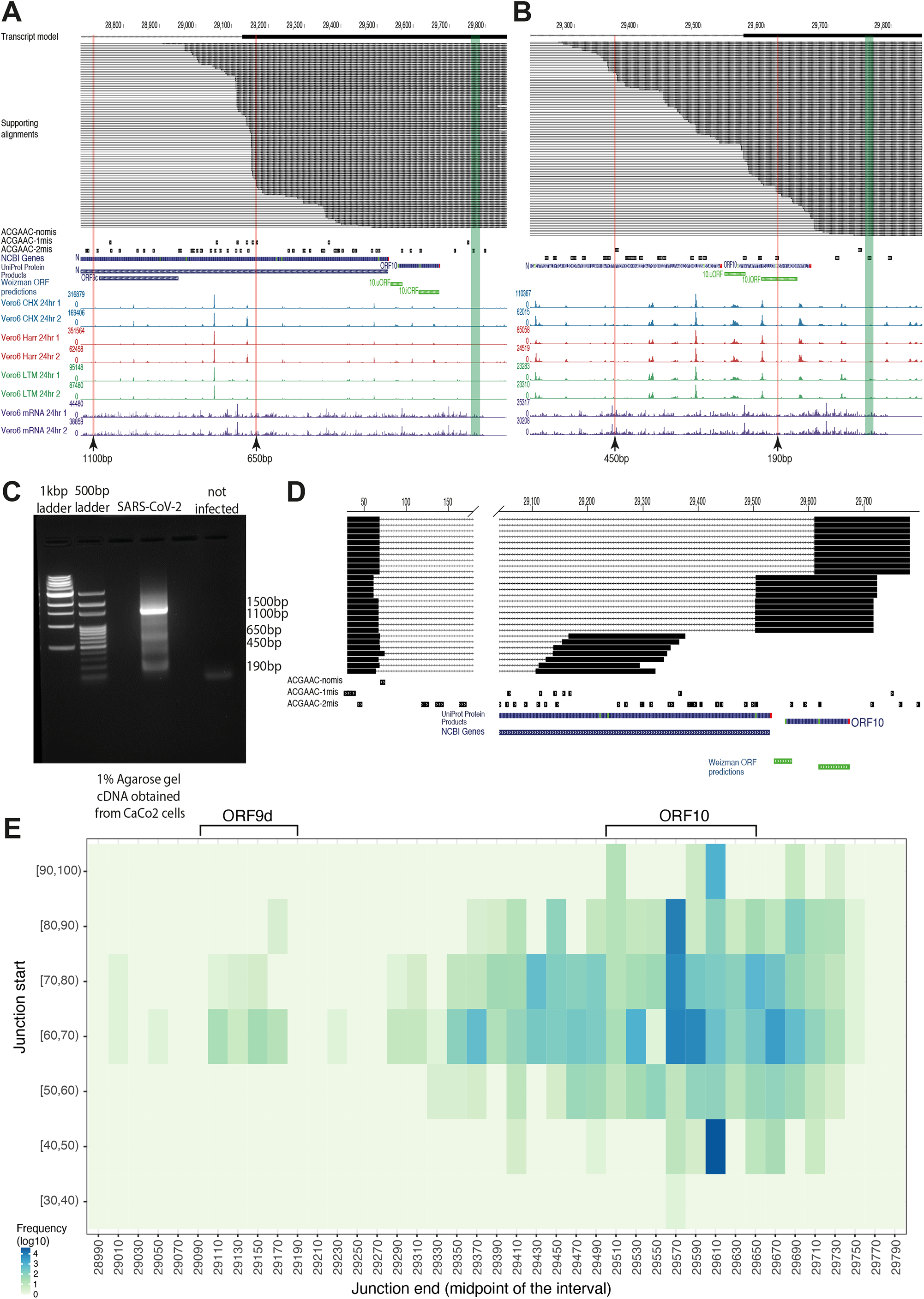
Independent sgRNAs encode ORF9d and ORF10. (**A-B**) UCSC Genome Browser track showing the alignments of NRCeq reads assigned to ORF9d (**A**) and ORF10 (**B**). The figure also includes tracks showing the location of TRS-B sequences with 0,1 or 2 mismatches, NCBI Genes, Uniprot Protein Products, ORF predictions and ribosome footprints^19^. Arrows indicate the genomic position of the products found in the bands of the gel in (**C**). (**C**) Agarose gel electrophoresis after PCR amplification of short sgRNA species. The band at 1500nt, 650nt and 190nt correspond to the expected size for the amplicons of full-length N ORF9d and ORF10 respectively. The bands at 450 and 1100 did not correspond to the size of any assembled transcript models. (**D**) Representative alignments of Illumina DNA sequencing data (250ntx2) of short, PCR-amplified sgRNAs. The bands at 1500nt, 650nt, 450nt and 190nt from the gel in (**C**) were purified and sequenced. (**E**) Heatmap showing the location and abundance of split alignments connecting the viral 5′UTR with downstream regions. The figure was generated using the Illumina DNA sequencing data as in (**D**). The y-axis reports the genomic coordinate upstream of the junction, whereas the x-axis reports the genomic coordinate downstream of the junction. The color scale reports the number of reads that support each junction (log10 transformed) after binning the genome in intervals of 10nt (x-axis) or 20nt (y-axis, the midpoint of every interval is displayed).

In addition, the NRCeq data also detected a novel canonical and capped sub-genomic RNA internal to, and in-frame with, the N protein. In line with the nomenclature of other ORFs internal to N, we named this sgRNA ORF9d. ORF9d is supported by 107 reads with junctions between the canonical TRS-L and non-canonical TRS-B sequences located at genomic position 29,128. Similarly to what we observed for ORF10, we also found ribosomal footprinting peaks supporting the translation of ORF9d (**Fig. 3A**). This ORF encodes the last 103 C-terminal amino acids of the N protein, which correspond to its C Terminal Domain (CTD), an important regulatory domain with RNA binding capacity^20^. We next sought to confirm the existence of sgRNAs encoding ORF10 and ORF9d using an orthogonal technique independent of Nanopore sequencing. To this end we designed a set of primers in a region shared by every sgRNA and performed RT-PCR with a short extension time, in order to favour amplification of short RNAs over longer, more abundant overlapping transcripts. We detected amplicons in the expected size ranges of both ORF10 and ORF9d (∼190bp and ∼690bp respectively, **Fig 3C**), which we purified and sequenced by Illumina DNA sequencing (2×250bp read length). After aligning reads to the viral reference genome we confirmed the leader-to-body fusion events for both ORF9d (88 reads spanning the fusion) and ORF10 (1841 reads spanning the fusion, **Fig 3D,E**). Interestingly, these data also confirm the existence of a short isoform of ORF10 with a leader-to-body fusion internal to the ORF, as previously reported^19^.

### Expression of non-canonical sgRNAs

Among the 21 sgRNA models obtained through NRCeq, 7 were non-canonical transcript models that lack one or more of the canonical features described above. In particular, we detected transcript models that either do not terminate at the 3′ of the reference genome or that are 5′ incomplete, i.e. transcribed from a single, contiguous genomic region and thus lacking the typical TRS-L/TRS-B fusion. The alignment of canonical sgRNAs to the viral reference genome consist of three regions: a short region (∼70nt) that aligns to the 5′ of the genome, a large gap (∼21-29kbp) and long region aligning to the 3′ end of the genome. We reasoned that a high number of mismatches in the 5′ region might cause the aligner to prefer a 5′ truncated alignment (i.e. with 5′ soft-clipped bases) rather than opening a large gap (even when using splice aware mode), thus creating artifactual support for 5′ truncated non-canonical transcript models. To exclude this possibility, we manually inspected the alignments supporting non-canonical transcript models (**Suppl. Fig. 6** and **Suppl. Information**). When looking at all reads, we found that a high percentage of alignments (38.8%) from the standard DRS library had between 1 and 10 softclipped nucleotides at the 5′ end (**Suppl. Fig. 7A**), likely reflecting alignment mismatches caused by poor basecalling accuracy near read ends^13^. On the other hand, NRCeq alignments had between 10 and 20 softclipped nucleotides at the 5′ end, likely due to incomplete adapter trimming (**Suppl. Fig. 7B-C** and **Suppl. Information**). Surprisingly, when we repeated this measurement using alignments that support non-canonical transcripts, we found an increase in soft-clipping length (**Suppl. Fig. 7E and Suppl. Information)**, which ranged between 0 and 80nt. In particular, two related groups of non-canonical transcript models (groups #2 and #3, see **Suppl. Fig. 6** and *non-canonical transcript model classification* in **Suppl. Information**) showed a high number of reads with a 5′ soft-clipping between 50 to 80nt (**Suppl. Fig. 7E and Suppl. Information**). These results suggest that reads supporting non-canonical sgRNAs could in reality derive from canonical sgRNAs, but due to the high error rate their 5′ leader sequence can not be mapped properly. These artefacts inflate the expression estimates of non-canonical sgRNAs and should be carefully evaluated when analysing Nanopore data for complex, nested transcriptomes (examples in **Suppl. Information** and **Suppl. Fig. 8**). However, despite these artefacts, we were able to find a small number of genuine alignments where the NRCeq adapter is directly linked to the sgRNA body (**Suppl. Fig. 9**). Altogether, these data demonstrate the existence of non-canonical sgRNAs that possess a 5′ cap but lack a 5′ leader sequence, although their expression level is low and artificially inflated by mapping errors.

Finally, we inspected the two deletions found in transcript models encoding ORFM and N. The deletion in the transcript model which encodes for M maintains the reading frame while the deletion in N causes a frameshift (**Suppl. Fig. 10 A-B-C**). The existence of these deletions has been confirmed by Illumina data (see **Materials and Methods**) and they are not specific for particular cell types or viral strains, as alignments supporting them have been observed in all NRCeq and DRS dataset (data not shown).

### The NRCeq data correctly quantifies annotated sgRNAs

After assembling the SARS-CoV-2 transcriptome we quantified the expression of sgRNAs using the full-length NRCeq reads as well as reads from the standard DRS protocol. Reads were aligned to the assembled transcriptome and quantified using NanoCount^21^. We then binned sgRNAs by length and calculated the relative expression in order to compare Nanopore data with reference expression values obtained by Northern Blot^22^ (**Fig. 2C** and **Suppl. Fig. 11**). We found that NRCeq and standard DRS provide expression estimates that are consistent with Northern Blot results (Spearman correlation coefficients of 0.998 and 0.928 for DRS and NRCeq respectively. See **Suppl. Table 4**). Specifically, both Nanopore sequencing data and Northern Blot analysis show that the expression of sgRNAs increases from longer to shorter transcripts. The main discrepancy between expression estimates based on NRCeq and Northern Blot is in the NCAP sgRNAs, which appear more abundant in NRCeq. This is likely a quantitative bias introduced by the fact that shorter RNAs are more likely to be sequenced full-length. In line with this, we also observed that NRCeq understimates the expression of spike ORF, which is encoded in the longest sgRNA. In contrast, this bias was absent when quantifying sgRNA expression using standard DRS data (**Suppl. Fig. 11**).

### The expression level of sgRNAs is conserved between different cell lines

To quantify the viral transcriptome across diverse samples, we performed a quantification analysis on individual DRS SARS-CoV-2 datasets (**Suppl. Table 1**). These datasets were derived from three different cell lines (CaCo2, Vero and CaLu3) infected with three different viral isolates (**Suppl. Table 1**). The sequencing throughput for each sample was similar and ranged between 692,530 and 1,588,319 reads. To assess the viral load for each dataset, we aligned reads to the reference SARS-CoV-2 genome and found that on average 47.8% aligned to the viral reference (**Fig. 4A-B)**. However, samples derived from CaCo2 cells had a significantly lower fraction of reads that aligned to the reference compared to those from Vero cells (CaCo2 26.7%, Vero 57.8%) or from CaLu3 cells (61.7%). This observation agrees with previous reports documenting lower viral titers in CaCo2 cells^17^.

**Figure 4.**
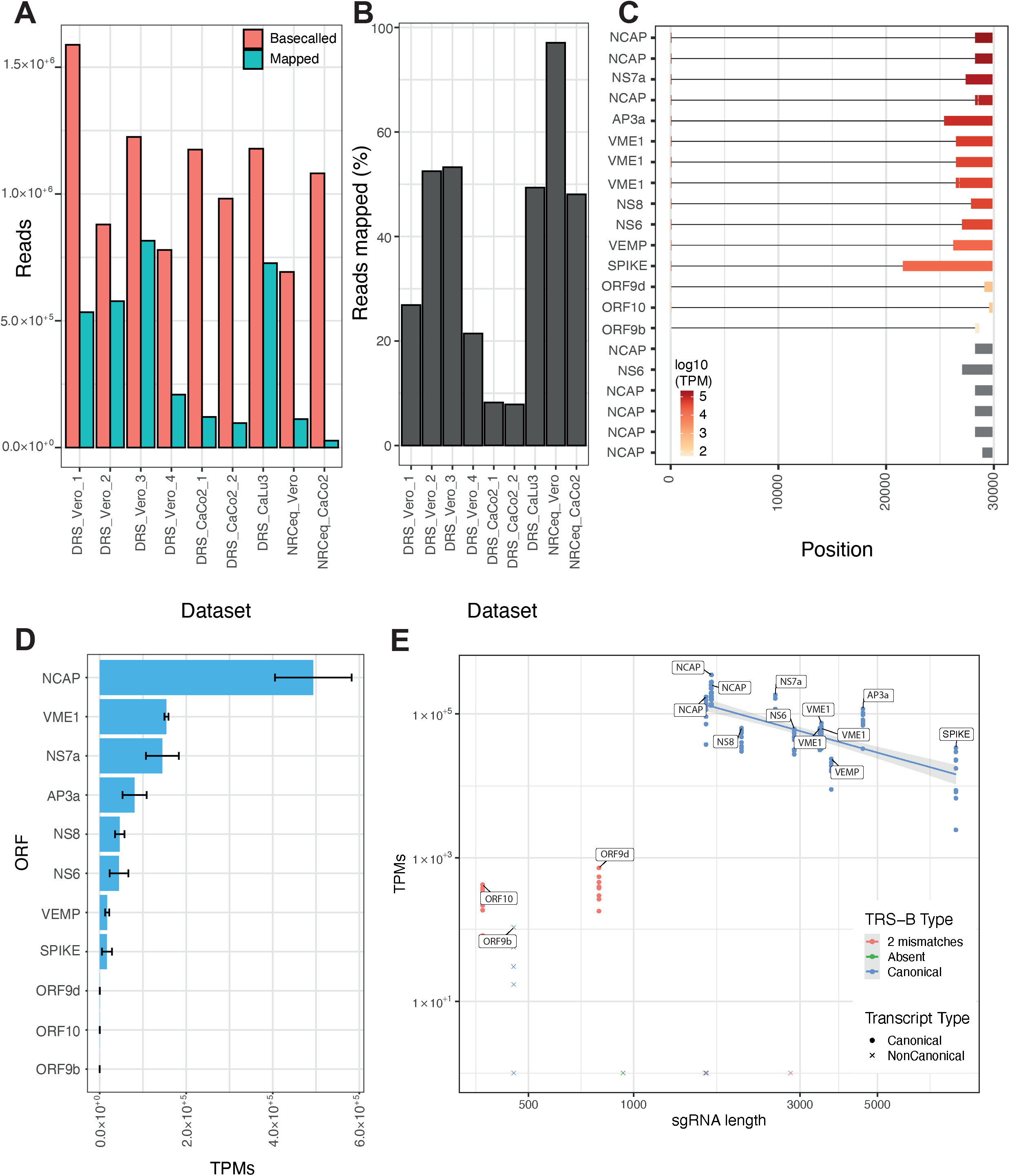
Expression profiling of sgRNAs in different cell lines. (**A-B**) Number (**A**) and percentage (**B**) of reads basecalled and mapped to the SARS-CoV-2 genome for each dataset. For the NRCeq datasets, the percentage is calculated as a fraction of the total number of capped reads. (**C**) Structure and mean expression of sgRNA transcript models across all datasets. The labels on the y-axis refer to the first ORF identified in each sgRNA. (**D**) Cumulative expression of all transcripts that code for each ORF. The values correspond to the mean expression of ORFs across all samples. The error bars report the combined standard deviation of the expression of the transcript models encoding each ORF (see **Materials and Methods**). (**E**) Scatter plot showing TPMs versus transcript length in each dataset.

We then quantified the sgRNA expression in each dataset by mapping the reads to the NRCeq assembly and expression estimates with NanoCount^21^ after excluding incomplete sequencing reads or RNA degradation products through a full-length filtering strategy (**Suppl. Information** and **Suppl. Fig. 12**). Despite different viral loads among samples, we observed that relative sgRNA expression was consistent between cell lines as well as viral strains, with canonical sgRNAs expressed at significantly higher levels than non-canonical ones (**Fig. 4C-D** and **Suppl. Fig. 13**). Additionally, we confirmed in all samples that the expression of canonical sgRNAs is negatively correlated with their length (**Fig. 4E**; Pearson correlation coefficient -0.71, p-value=4.9×10^−15^). Unsurprisingly, ORF9d and ORF10 had relatively low expression despite their shorter length, likely due to their non-canonical TRS-B sequence.

## Discussion

In this work we have applied a recently developed technique, called Nanopore ReCappable Sequencing (NRCeq), to profile the repertoire of full-length, capped RNAs produced by the SARS-CoV2 virus.

We first generated an annotation of capping sites across the viral genome, in the form of genome browser tracks, which allow determining the RNA start sites without relying on a transcriptome assembly. These data, which are the first of their kind, provide a valuable resource for the scientific community seeking to better explore the transcriptional mechanisms of SARS-CoV2.

We also leveraged the NRCeq data to assemble a *de novo* SARS-CoV-2 transcriptome. We could identify transcript models supporting all canonical sgRNAs with the exception of ORF7b, which is thought to be expressed from the ORF7a mRNA by leaky ribosome scanning. The absence of a transcript model for this ORF in our dataset could be explained by the lower sequencing depth of NRCeq or by biological differences in the expression (or precise location) of this sgRNA, as previously suggested by others^23^.

Our data support the expression of an sgRNA for ORF10, whose existence was recently debated^4,8,19^. Using NRCeq we could reliably identify over 100 capped reads that span the leader-to-body junction of ORF10 and have TRS-B sequences with one or two mismatches located upstream of the start codon. Our data also provide evidence for the existence of a novel canonical sgRNA that encodes a truncated version of the N protein, which we named ORF9d. Together with ORF9b and ORF9c, this is the third known ORF internal to N, but they are all of unknown functional significance. Although direct evidence that ORF9d produces a functional protein is still lacking, it might have a direct regulatory role on the abundance of sgRNAs, since the polypeptide that it encodes overlaps the C-terminal region of the N protein, which was recently shown to specifically interact with TRS sequences^24^. Alternatively, it is possible to speculate that ORF9d could have a function independent of its coding potential or act as substrate for the evolution of new viral proteins. This hypothesis is supported - at least in part - by the presence of multiple ORFs with alternative start codons internal to ORF9d, the longest of which encodes a polypeptide of 36 amino acids.

In addition to canonical sgRNAs, our assembly also identified non-canonical ones that either lack the leader-to-body fusion or terminate upstream of the genomically encoded polyA tail. Through the careful examination of the alignments, we observed that the expression of these sgRNAs was artefactually inflated by mapping errors. This observation stresses the importance of carefully optimising mapping parameters when dealing with reads arising from nested transcripts. Despite these technical biases, we still detected a number of properly mapped reads that support the expression of 5′ truncated non-canonical sgRNAs. These RNAs are capped and contain intact ORFs, and are therefore potentially protein coding. The functional significance of not having the 5′UTR region provided by the leader-to-body fusion is still unclear, but it might have regulatory effects, for example modulating RNA-protein interactions or altering RNA structure or stability.

We also quantified the expression of sgRNAs across sequencing datasets, finding that expression estimates obtained by both NRCeq and standard RNA-Seq are in line with the expectations from northern blot experiments. We also observed that the expression levels are remarkably stable across cell lines and viral isolates, suggesting the presence of a robust mechanism that regulates RTC activity. In fact, in line with previous observations^4^, we observed that the sgRNA expression levels are inversely proportional to their length. This phenomenon can be explained by the fact that the RTC has a certain probability of switching templates at each TRS-B sequence that it encounters; longer negative sense intermediates - having a higher number of internal TRS-B sequences - are less likely to be transcribed in their entirety. Our data shows that length alone explains 50.0% of the variance in sgRNA expression. It is plausible that part of the remaining variability is due to other factors, such as the TRS-B sequence itself. In line with this hypothesis, we found that short sgRNAs lacking a canonical TRS-B sequence (i.e. ORF9d and ORF10) have a much lower expression level than expected for their length.

In conclusion, NRCeq is a robust technique that permits assembly and quantification of complex transcriptomes. By providing an annotation of capping sites and annotating novel, capped transcripts, our work helps to shed light on the complex mechanisms that regulate SARS-CoV-2 transcription and sgRNA formation.

## Materials and Methods

### Cell culture and samples preparation

#### CaCo2 (sample 2) and CaLu3 samples for DRS

##### Cells and virus

Vero E6 cells (Vero C1008; clone E6–CRL-1586; ATCC) were cultured in Dulbecco’s modified Eagle medium (DMEM) supplemented with nonessential amino acids (NEAA, 1x), penicillin/streptomycin (P/S, 100 U/mL), HEPES buffer (10 mM) and 10% (v/v) Fetal bovine serum (FBS). CaCo2 (Human epithelial colorectal adenocarcinoma cells, ATCC HTB-37) cells were cultured in Minimum Essential Medium (MEM) supplemented with NEAA (1x), P/S (100 U/mL), HEPES buffer (10 mM), sodium pyruvate (1 mM), and 20% (v/v) FBS. CaLu3 (Human lung cancer cell line, ATCC HTB-55) were cultured in MEM supplemented with NEAA (1x), P/S (100 U/mL), HEPES buffer (10 mM), sodium pyruvate (1 mM) and 10% (v/v) FBS. A clinical isolate hCoV-19/Italy/UniSR1/2020 (GISAID accession ID: EPI_ISL_413489) was isolated and propagated in Vero E6 cells. All the infection experiments were performed in a biosafety level-3 (BLS-3) laboratory of Microbiology and Virology at Vita-Salute San Raffaele University, Milan, Italy.

##### Virus Isolation

An aliquot (0.8 mL) of the transport medium of the nasopharyngeal swab (COPAN’s kit UTM® universal viral transport medium—COPAN) of a mildly symptomatic SARS-CoV-2 infected patient was mixed with an equal volume of DMEM without FBS and supplemented with double concentration of P/S and Amphotericin B. The mixture was added to 80% confluent Vero E6 cells seeded into a 25 cm^2^ tissue culture flask. After 1 h adsorption at 37°C, 3 mL of DMEM supplemented with 2% FBS and Amphotericin B were added. After five days, cells and supernatant were collected, aliquoted, and stored at −80°C (P1). For secondary (P2) virus stock, Vero E6 cells seeded into 25 cm^2^ tissue culture flasks were infected with 0.5 mL of P1 stored aliquot, and infected cells and supernatant were collected 48 hours post-infection and stored at −80° C. For tertiary (P3) virus stock, Vero E6 cells were infected with 0.2 mL of P2 stored aliquot and prepared as above described.

##### Virus Titration

P3 virus stocks were titrated using both Plaque Reduction Assay (PRA, PFU/mL) and Endpoint Dilution Assay (EDA, TCID_50_/mL). For PRA, confluent monolayers of Vero E6 cells were infected with 10-fold-dilutions of virus stock. After 1 h of adsorption at 37°C, the cell-free virus was removed. Cells were then incubated for 46 h in DMEM containing 2% FBS and 0.5% agarose. Cells were fixed and stained, and viral plaques were counted. For EDA, Vero E6 cells (3 × 10^5^ cells/mL) were seeded into 96 wells plates and infected with base 10 dilutions of virus stock. After 1 h of adsorption at 37°C, the cell-free virus was removed, and complete medium was added to cells. After 48 h, cells were observed to evaluate CPE. TCID_50_/mL was calculated according to the Reed–Muench method.

##### Infection experiments

Caco2 and CaLu3 cells were seeded on 25 cm^2^ tissue culture flasks until 80% confluency. Then, flasks were infected with SARS-CoV-2 at 0.1 multiplicity of infection (MOI). After 1 h of virus adsorption, cells were washed with PBS, and further cultured at 37°C for 48 h with 4% and 2% FBS, respectively. After a PBS wash, enzymatic dissociation was performed for 4–6 minutes at 37°C in 1 mL TrypLE (Invitrogen), then cell pellets were washed with ice-cold PBS and lysed with 1 mL of TRIzol (Invitrogen). The samples were stored at -80° C for subsequent RNA extraction.

##### RNA extraction and Nanopore direct RNA sequencing

RNA was isolated using Trizol-chloroform extraction followed by purification using RNeasy Mini kit with Dnase treatment (Qiagen) according to the manufacturer’s protocol.

Standard Nanopore DRS libraries were prepared from 4ug of total RNA from infected cells using the SQK-RNA002 kit and following the standard protocol with the same adaptation as in Kim et al.^4^ Sequencing was done on a FLO-MIN106 flowcell on a GridION instrument.

#### CaCo2 (sample 1) and Vero (sample 3) samples for DRS

Isolation and growth of SARS-CoV-2/human/Liverpool/REMRQ0001/2020 has been previously described^25^. Briefly, confluent CaCo2 or VeroE6 cells seeded in a 75cm2 flask were infected with a multiplicity of infection of 0.1 and total RNA was harvested using Trizol reagent as previously described^8^ after 24 hours infection. Briefly we extracted total RNA as per the manufacturer’s protocol except we wash-precipitated RNA three times in 1ml 75% ethanol, we processed the RNA immediately and poly(A) selection was done using Dynabeads. We performed the RNA extraction, polyA selection, recapping and DRS without any pause or storage. All experiments with live virus were performed at the BSL3 facility within the School of Medical Sciences at the University of Bristol, UK.

#### NRCeq experiments (CaCo2 and Vero cells)

Viral RNA was decapped and recapped as previously described^13,26,27^. Briefly, 1.5-6ug of poly(A) selected RNA was decapped using 1.5 μL yDcps (NEB, #M0463) in 1X yDcpS reaction buffer (10 mM Bis-Tris-HCl pH 6.5, 1 mM EDTA) in 50 μL total volume for 1 h at 37 ºC. The decapped RNA was purified using an RNA Clean and Concentrator (Zymo Research, #R1013) using the manufacturer’s recommended protocol and eluted in 30 μL of RNase-free water. The decapped RNA was Recapped with 6 μL Vaccinia Capping Enzyme (VCE) (NEB, #M2080) in 1X VCE reaction buffer (50 mM Tris HCl, 5 mM KCl, 1 mM MgCl2, 1 mM DTT, pH 8), 6 μL E. coli Inorganic Pyrophosphatase (NEB, #M0361), 0.5 mM 3′-azido-ddGTP (Trilink, #N-4008), 0.2 mM S-adenosylmethionine (SAM) (NEB, #B9003) in 60 μL total volume for 30 min at 37 ºC. The recapped RNA was purified with RNA Clean and Concentrator as above.

The azido-ddGTP recapped RNA (1-2 μg) was concentrated to approximately 7 μL using a SpeedVac vacuum concentrator (Savant). Copper-free Click Chemistry reactions were performed in a total volume of 50 μL, containing 25% v/v PEG 8000 (NEB, #B1004) and 20% v/v acetonitrile (Sigma-Aldrich, #271004) in 0.1 M sodium acetate buffer, pH 4 (10X, Alfa Aesar, #J60104) and 10 mM EDTA (50x, Invitrogen, #15575-038). Azido-ddGTP recapped RNA and the 3′-DBCO RNA adapter (200 nmol, final concentration of 4 μM) were added and shaken for 2 h at room temperature. Then, acetonitrile was removed by brief concentration on a SpeedVac, and the adapted RNA was purified using RNA Clean & Concentrator (Zymo Research, #R1013) following the protocol to separate large RNA (desired) from small RNA (excess adapter).

## Bioinformatic analyses

### Viral reference genome

The reference viral genome fasta was downloaded from the UCSC Genome Browser^28^ and modified according to the criteria proposed by Kim et al^4^.

### Basecalling

In-house sequenced datasets were basecalled with Guppy, available to ONT customers via their community site (https://community.nanoporetech.com) (version in **Suppl. Table 1**). Quality controls, including information on the quality of the reads and on the outcome of the basecalling, were performed through pycoQC^29^ (v.2.5.2).

### NRCeq data processing

#### Trimming of the NRCeq adapter

The 5′ end adapter was identified and trimmed from ReCapped read fastq files using Porechop, as described in Mulroney et al.^13^. A subsequence of the adapter, TCCCTACACGACGCTCTTCCGA, was added to the Porechop adapters file and Porechop was executed with the following parameters:

--barcode_diff 1

--barcode_threshold 74

#### Mapping to the SARS-CoV-2 genome

Trimmed ReCapped reads were mapped against the reference viral genome with minimap2^18^ (v2.17-r974) using the same parameters as those used by Kim et al^4^. For the analysis on the impact of different minimap2 parameters on the amount of soft-clipping at the 5′ end of NRCeq alignments (in **Suppl. Information**), three parameters combination were used: standard DRS long-read conditions (as in minimap2^18^), splice-aware conditions (as in minimap2^18^) and Kim et al.^4^ conditions.

#### Reads statistics

The number of Basecalled and Trimmed reads was obtained counting the entries of the fastq files. The number of Mapped Reads was calculated through samtools view^30^ (v1.10-76-g65c8721). Only primary alignments have been kept into account (using the flags -F 2324 for positive strand alignments, -F 2308 -f 16 for negative strand alignments).

The total coverage of the viral genome and of the 5′ and 3′ ends was calculated through bedtools genomecov^31^ (v2.27.1) using the -split, -5 or -3 flags for total, 5′ or 3′ coverage respectively.

Tracks for UCSC Genome Browser were generated in bedgraph format through bedtools^31^ genomecov using the following parameters:

-ibam <input bamfile>

-bg

-trackline

-split

-strand +/-

-5/3 (for 5′ or 3′ ends)

#### Peak calling

A peak calling analysis was performed on the coverage of the 5′ ends of NRCeq positive strand alignments to establish the distribution of the alignment start sites per genomic position. Briefly, the coverage per genomic position was grouped in rolling windows of 10 nucleotides through the function zoo::rollapply^32^ and the partial sum for each interval was calculated. Finally the peak calling was performed through the function ggpmisc::stat_peaks^33^ (parameters: ignore-threshold = 0.0005 and span = 5). Results were plotted through tidyverse::ggplot2^34^. A bed file with NRCeq peaks is displayed in **Suppl. Table 2**.

#### Transcriptome assembly

In order to build a NRCeq assembly of the viral transcriptome, NRCeq primary alignments to the reference genome were sorted and indexed through samtools^30^. Transcriptome assembly was performed with Pinfish with the following parameters:

-spliced_bam2gff -s <NRCeq bam file> (converts the input bam file in a GFF)

-cluster_gff -c 100 -d 80 -e 100 -p -1 -a <clusters.tsv gff file> (cluster reads in the gff)

-polish_clusters -f -a <clusters.tsv> -c 100 -o <out fasta file> <NRCeq bam file> (polishes the clusters and outputs the consensus fasta file)

Pinfish parameters above were obtained by manual tuning in order to obtain the best possible assembly, balancing the number of redundant transcript models per sgRNA and a sufficient coverage of the isoforms present in the assembly.

The consensus fasta file obtained was then mapped to the reference genome with minimap2^18^, using the parameters in the section above. Finally the alignment file was converted to bed with bedtools^31^ bamtobed -bed12.

#### Assignment of ORFs to transcript models

To classify sgRNAs based on the protein that they encode, we used orf-annotate^35^, a tool that first identifies the first ORF in a sequence then translates it and aligns the resulting amino acid sequence to the reference proteome from Uniprot^14^. orf-annotate was run with the following parameters: --bedfile <assembly bed file> --fasta < extracted fasta sequence> --proteins-fasta <reference proteome fasta>.

Two deletions were present in two different transcript models of the NRCeq assembly. Their genomic coordinates were extracted from the bed file of the assembly using bedparse introns^36^.

In order to assess if these deletions were real biological entities or mapping artefacts, we analysed Illumina datasets obtained from the infection of Vero and CaCo2 cells by Liverpool viral strain in Matthews Lab.

#### Deletion analysis using Illumina data

Illumina data from DRS_Vero_3 and DRS_CaCo2_1 datasets (Matthews Lab) were mapped to the reference genome fasta with STAR^37^ (v 2.7.9a) with the following commands:

> STAR --runMode genomeGenerate --genomeDir <directory for genome indexes> --genomeFastaFiles <reference genome fasta> --genomeSAindexNbases 7 (to build genome index for short genomes)
>
> STAR --runThreadN 8 --genomeDir <directory for genome indexes> --readFilesIn <fastq Illumina files> --outFilterIntronStrands None --outSJfilterOverhangMin 10 12 12 12 --outFileNamePrefix <prefix for output files> (to map PE reads)

The resulting alignment file was filtered (samtools -F 2316) and converted to a bed file (bedtools^31^ bamtobed -bed12).

The gaps of each alignment were extracted from the bam file through bedparse^36^ introns (v0.2.3) and saved in bed format. Similarly, we used bedparse introns to extract the coordinates of the deletions from the NRCeq transcript models.

To assess if gaps recovered from Illumina alignments supported the existence of the deletions observed in two transcript models of the NRCeq assembly, we used the command bedtools^31^ intersect with options -f 0.8 -r -a <bed file of the NRCEq assembly deletions> -b <bed file of introns from Illumna alignments> -wb. This command gives the intersection between the genomic coordinates of the deletions with the introns of the alignments of the Illumina data. Once obtained the overlapping sequences and the corresponding id of Illumina alignment, we extracted the alignments overlapping the deletions from the sorted and indexed Illumina bam file through a custom python script.

We then converted these alignments in bed format through bedtools^31^ bamtobed -bed12, uploaded the bed file in the UCSC Genome Browser and manually inspected the alignments supporting the deletions.

### DRS data processing

Mapping and calculation of standard statistics (number of reads and coverage profiles) were done as already described for the NRCeq data.

Peak calling was done as described for the NRCeq data, but the ggpmisc::stat_peaks^33^ function was called with the parameter span=40.

### NRCeq and standard DRS dataset quantification

#### Mapping to the NRCeq assembly

NRCeq and standard DRS datasets were mapped to the NRCeq assembly in order to quantify the transcriptional landscape of SARS-CoV-2. Fastq files were mapped to the assembly extracted fasta file through minimap2 with the following parameters:

-t 16

-ax map-ont

-p 0

-N 10

#### Quantification

In order to quantify the expression of each transcript model, NRCeq and standard DRS alignment files were filtered keeping only primary and secondary alignments (samtools **-F** 2068), then they were sorted (samtools sort) and indexed (samtools index).

Two types of quantification against the NRCeq assembly were performed on these files: “all reads” quantification, that is the quantification of all the primary and secondary alignments, and the “full length” quantification, that is the quantification of the primary and secondary alignments having the 5′ end falling in an interval of 200 nucleotides centered at the first 5′ end nucleotide of each transcript model (see **Suppl. Information**).

The “all reads” quantification on the filtered and sorted datasets was performed through NanoCount^21^ with the following parameters:

-3 10

-5 10

-p align_score

-x

The “full-length” quantification was performed intersecting (samtools view -L) the filtered and sorted alignment files with a bed file with intervals of 200 nucleotides centered at the first 5′ end nucleotide of each transcript model. The resulting alignment files were quantified against the NRCeq assembly through NanoCount^21^ with the same parameters as above. Cumulative expression of ORFs were calculated by summing tpms of transcript models encoding the same ORF and calculating the mean across samples. Standard deviation for each ORF was calculated through the function combinevar in the R package *fishmethods*.

### ORF9d and ORF10 validation

PCR amplicons for short-read Illumina sequencing were produced in 50 μl of reaction buffer (10 mM dNTPs, 10 μM forward primer, 10 uM reverse primer, 3% DMSO, 25 units Phusion DNA polymerase (product number), and ∼500 ng template cDNA using 25 cycles (98C for 10 seconds, 65C for 10 seconds, 72C for 8 seconds) and a final 5 min extension at 72C. For primers we used the Artic protocol^38^ forward primer (annealing in the 5′UTR region, 5′–ACCAACCAACTTTCGATCTCTTGT–3′) and a reverse primer downstream of ORF10 (5′-CTCTCCCTAGCATTGTTCACTGTAC-3′). The template cDNA was generated from CaCo2 cells infected with SARS-CoV-2 or uninfected as a negative control. Distinct amplicons were purified from agarose gel slices using Qiagen gel extraction kit following the manufacturer’s instructions. The template cDNA was generated by retrotranscription of the RNA obtained from CaCo2 cells not infected or infected with SARS-CoV-2. Specifically, RNA was extracted using the miRNeasy mini kit (Qiagen, #1038703) and then retrotranscribed using the ImProm-II™ Reverse Transcription System (Promega, #A3801), according to the vendor’s instructions.

PCR products were separated on an agarose gel and the distinct amplicons were purified using the Qiagen gel extraction kit (#28706X4), following the manufacturer’s instructions. The gel purified amplicons were pooled in equimolar ratios and used as the input for RNA-Sequencing library preparation. Equimolar amounts of each band (1500, 1100, 650, 450 and 190bp) were pooled together to reach a total amount of 10ng. The amplicon DNA (1–10ng) was blunt-ended and phosphorylated, and a single ‘A’ nucleotide was added to the 3′ ends of the fragments in preparation for ligation to an adapter with a single-base ‘T’ overhang. The ligation products were then purified and accurately size-selected by agencourt AMPure XP beads. Purified DNA was finally PCR-amplified to enrich for fragments with adapters on both ends. All the steps above were performed on an automation instrument, Biomek FX by Beckman Coulter. The final purified product was quantitated prior to cluster generation on a Bioanalyzer 2100. The resulting library was sequenced for 250 bases in paired end mode on an Illumina MiSeq sequencer.

MiSeq adapters were trimmed using Reaper^39^ with the following parameters:

-geom no-bc -3pa GATCGGAAGAGCACACGTC ; for the R1 file

-geom no-bc -3pa CGGTGGTCGCCGTATCATT ; for the R2 file.

Reads were then aligned to the SARS-CoV-2 genome with STAR^37^ (v 2.7.9a) using the following parameters:

--outFilterIntronStrands None

--outSJfilterOverhangMin 10 12 12 12

Alignments were filtered with samtools^30^ (-F 2316) and only split alignments originating from the TRS-L were kept into account for further analysis.

The junction heatmap was built in R by counting the number of junctions that connected each genomic position bin in the TRS-L to a genomic position bin in the region upstream of ORF9d/10. The resulting numbers of junctions were then log10 transformed for visualisation purposes.

In order to confirm that adapted NRCeq reads fully aligned to the assembly, NRCeq untrimmed reads were aligned to the assembly fasta sequences preceded by the NRCeq adapter sequence using minimap2 with the following parameters:

-ax map-ont

-p 0

-N 10

Reads were filtered keeping only primary alignments (samtools -F 2324), then they were sorted (samtools sort) and indexed (samtools index). **Suppl. Fig. 5** reports such alignments as IGV tracks after filtering for less than 20 mismatches at the alignment start and mapping quality respectively higher than 0 (for ORF9d) or 30 (for ORF10).

### Soft-clipping analysis

In order to quantify the amount of the soft-clipping at the 5′ and 3′ ends of NRCeq and standard DRS alignments (against the reference genome), we used pysam to parse the CIGAR string of every alignment, annotating the amount of soft-clipping and hard-clipping at the 5′ and 3′ end. The soft-clipping amount for each dataset, respectively at the 5′ and 3′ end, was grouped in intervals and the results in TPMs plotted through an R script.

In the case of the peaks at 10-20 and 20-30 soft-clipped nucleotides at the 5′ end of alignments of NRCeq datasets (**Suppl. Fig. 7A**), alignments with respectively 10-20 and 20-30 soft-clipped nucleotides at the 5′ end were collected and grouped per transcription start sites. Normalised counts in NRCeq datasets were plotted per genomic position intervals (**Suppl. Fig. 7B-C**)

The same data processing was used to plot TPMs vs soft-clipping amounts, for NRCeq alignments supporting non-canonical transcript models, separated in categories as shown in **Suppl. Information**. These alignments were extracted from the NRCeq alignment files through a custom python script.

### Fast5 signal extraction

In some cases, we noticed alignments with a very high amount of soft-clipping at the 5′ end. To investigate these features, we used the python script in the section above to extract the soft-clipped sequences and we manually inspected them. We first BLAT^40^ the sequences against the viral reference genome and the human genome (hg38) and then extracted the raw ionic current data from the correspondent fast5 files. This was visualized using a custom matlab script.

### Northern Blot, NRCeq and standard DRS quantification comparison

Northern Blot quantification results from Ogando et al.^22^ were grouped in intervals of length 0.4 kp. NRCeq and Standard DRS counts obtained from Full-length quantification (see section above) associated with a specific transcript model, were assigned to the corresponding transcript model length and to the encoded ORF (see section above). TPMs were grouped per transcript length and divided in intervals of length 0.4 kp. Expression percent was calculated both for NRCeq and DRS datasets. Results were plotted through ggplot2^34^.

### Non-canonical transcript models analysis

In order to establish if non-canonical transcript models were supported by genuine capped reads, two types of analysis were performed.

To establish if the TRS-L had been softclipped, generating an artificial non-canonical alignment, the softclipped sequence at the 5′ of each NRCeq alignment to the viral genome which supported non-canonical transcript models was mapped to the 5′ UTR of the viral genome through parasail^41^ (v2.4.3) with the following parameters:

-a sw_trace

To prove the presence of capped non-canonical alignments, NRCeq untrimmed reads were aligned to each non-canonical transcript model sequence preceded by the adapter sequence using minimap2^18^ (same parameters used for the quantification).

## Data availability

The sequencing datasets generated in this study have been deposited at the European Nucleotide Archive (ENA) and are available under the id: PRJEB48830.

## Acknowledgements

We would like to thank the Genomics Unit of the Department of Experimental Oncology (European Institute of Oncology) and of the Center for Genomics Science (IIT) for their invaluable help with Nanopore sequencing runs. We also thank prof. Saverio Minucci (European Institute of Oncology) for his collaboration and the resources that he dedicated toward this project. We are also very grateful to Dr. Yuri D’Alessandria and John Buswell for providing useful reagents. We also thank New England Biolabs Inc. for their support of this research.

