## Supplementary Figures and Information for "Nanopore ReCappable Sequencing maps SARS-CoV-2 5′ capping sites and provides new insights into the structure of sgRNAs"

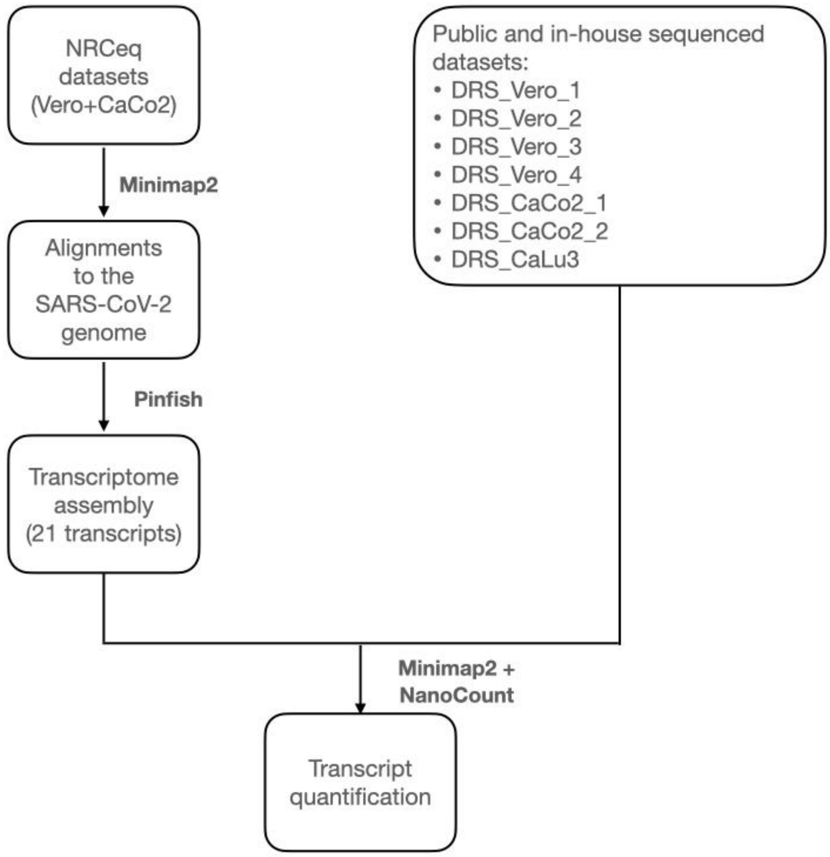


**Supplementary figure 1**. **Scheme of the pipeline.** (**A**) Schematic flow of the main pipeline to build the assembly and quantify SARS-CoV-2 transcripts.

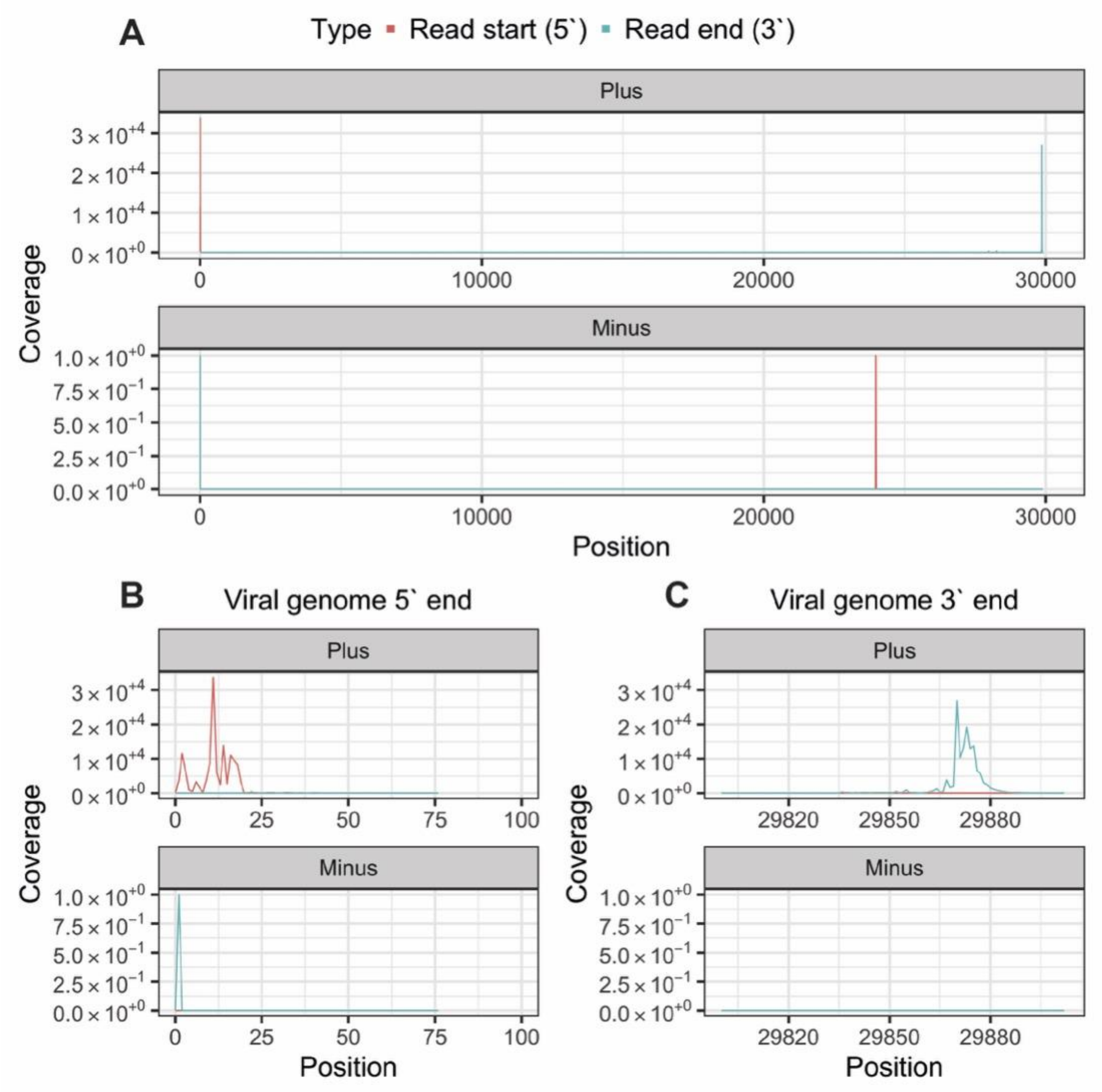


**Supplementary figure 2**. **NRCeq recovers full-length reads.** (**A**) Coverage of read starts and ends (5’ and 3’ coordinates) across the viral genome for the positive (top) and negative (bottom) strands. The y-axis shows the sum of the numbers of alignments covering each genomic locations in the two NRCeq datasets. (**C** and **B**) Same data as in A but magnified on the first (B) and last (C) 100nt of the viral genome.

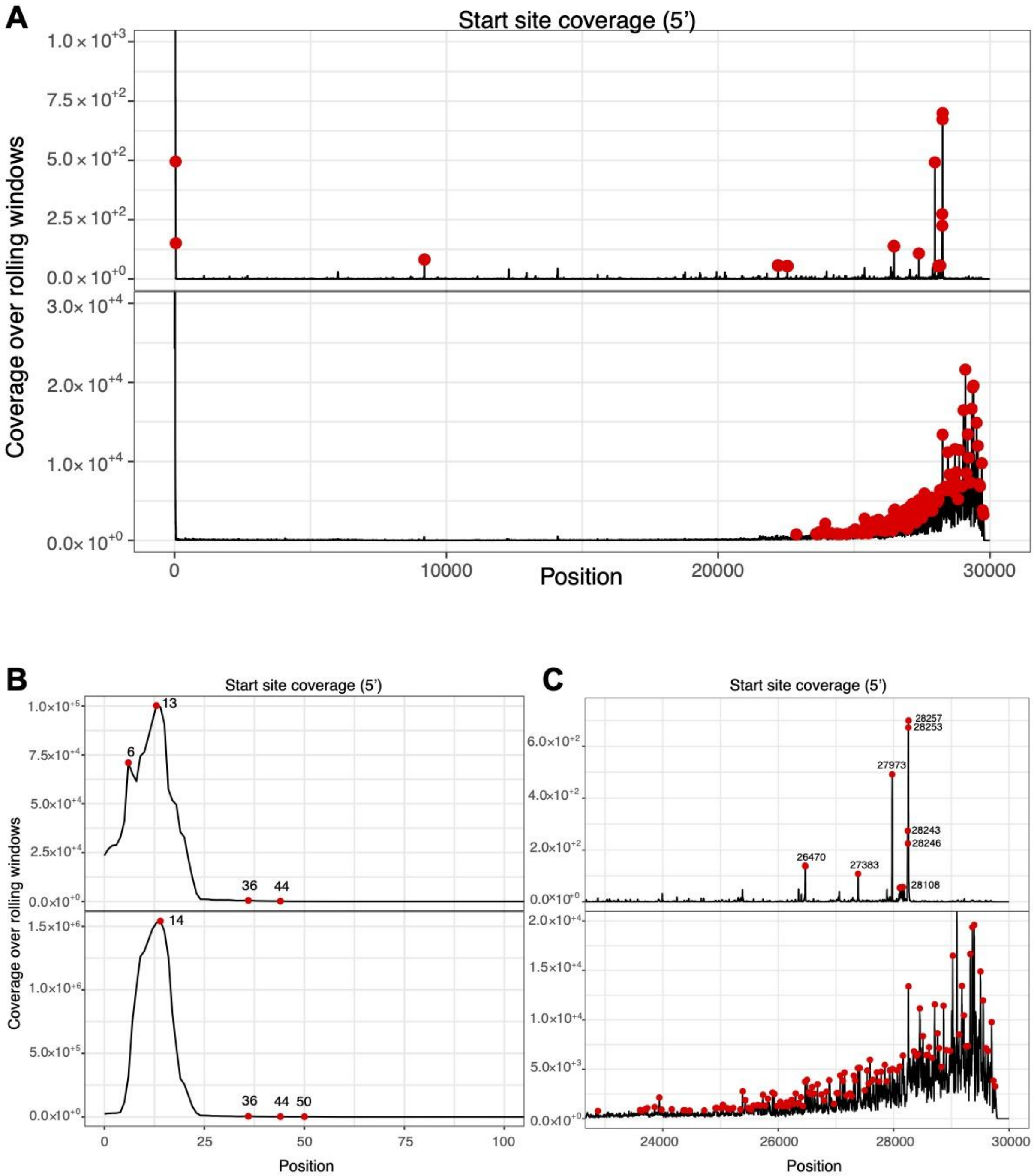


**Supplementary figure 3**. **Peak calling on NRCeq and Standard DRS data.** (**A**) Genome-wide peak calling of the coverage of the 5' of each alignment in NRCeq and Standard DRS datasets summed. Red points represent peaks called over a threshold by the algorithm (see **Materials and Methods**). (**B-C**) Same data as in A but magnified on the first (B) and last (C) 100nt of the viral genome.

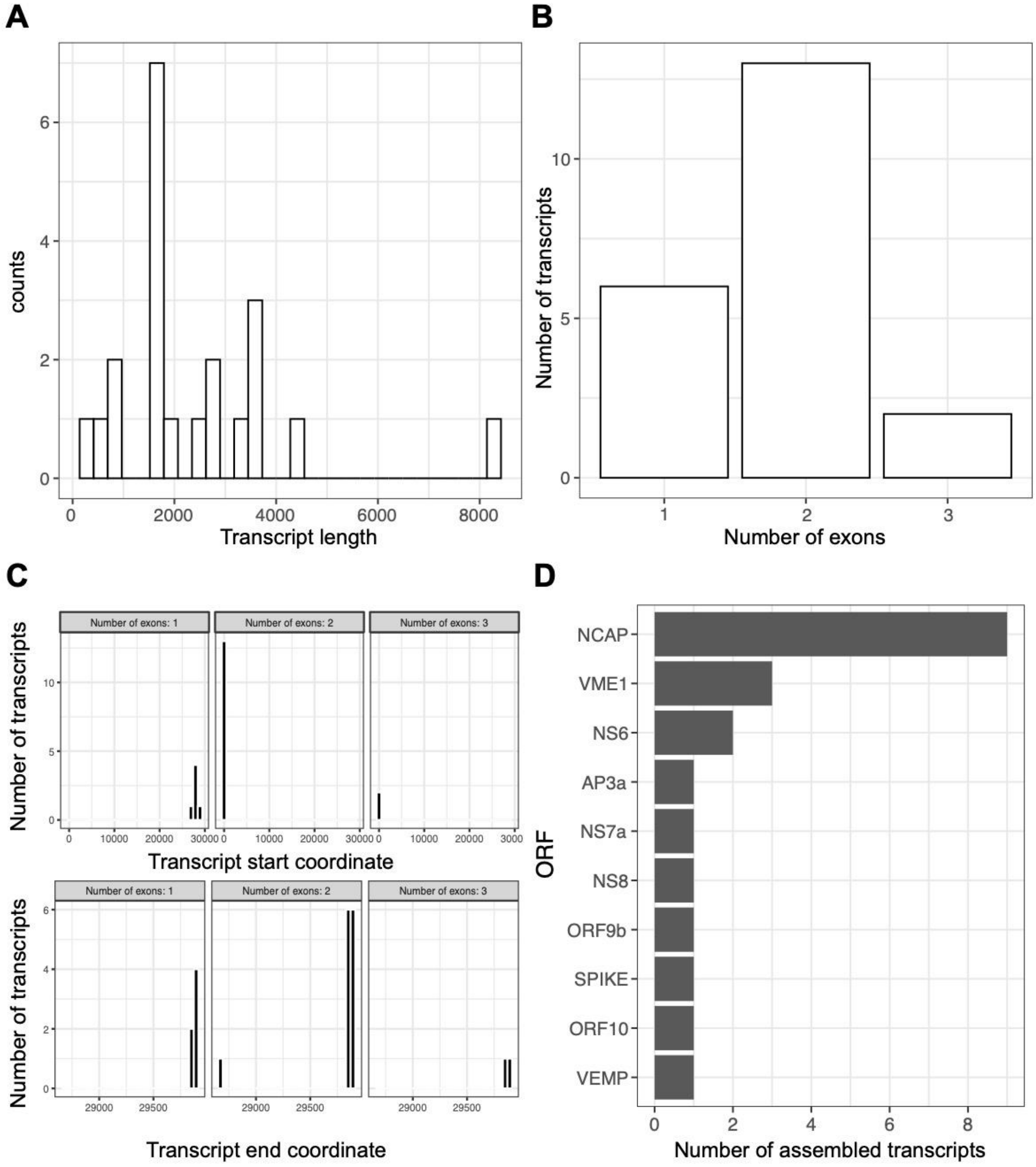


**Supplementary figure 4**. **Assembly quality controls.** (**A**) Size distribution of assembled transcripts. (**B**) Number of blocks per assembled transcripts. (**C**) Start and end coordinate of assembled transcripts. (**D**) Number of assembled transcripts that code for each ORF.

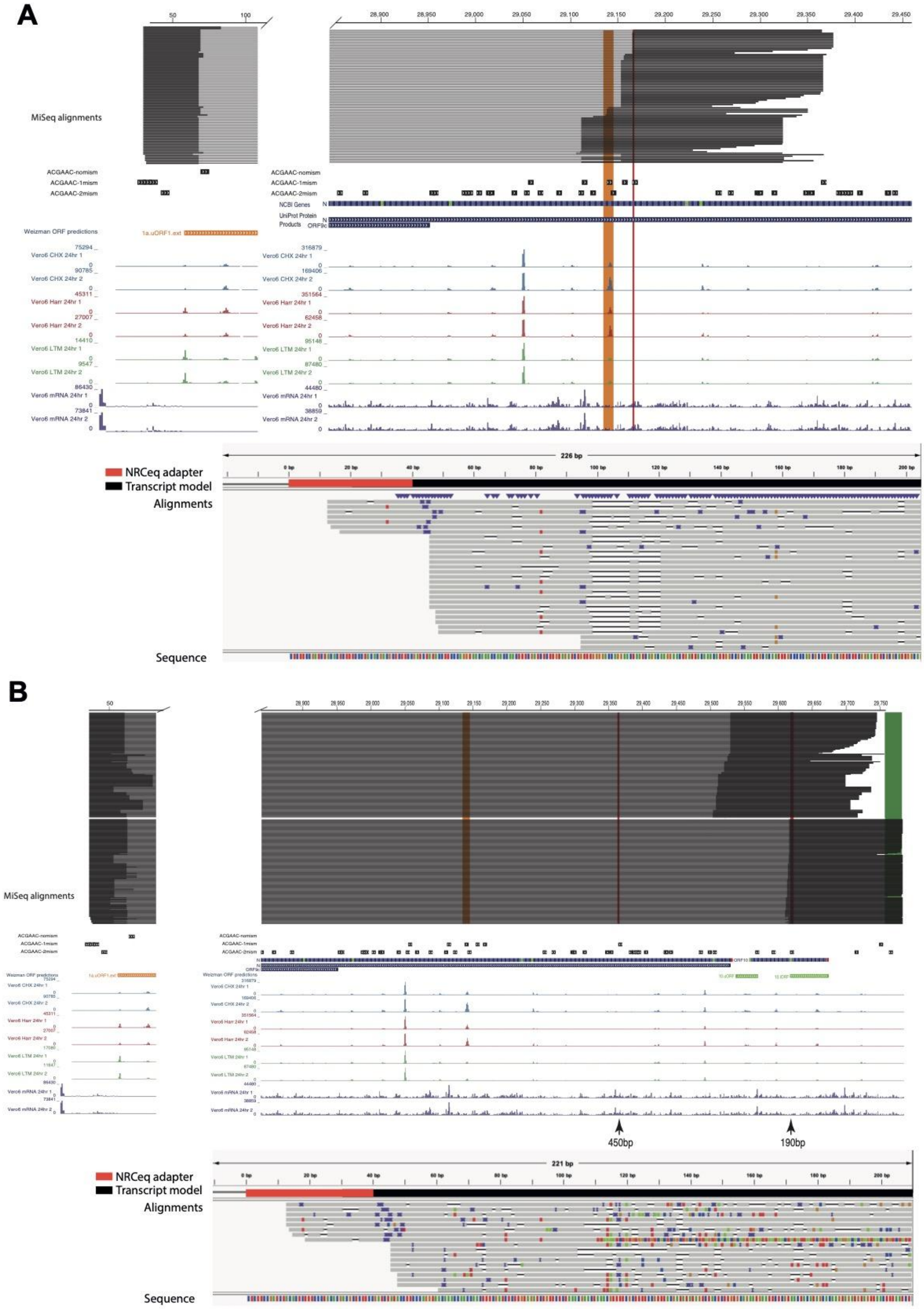


**Supplementary figure 5**. **ORF9d and ORF 10.** (**A-B**) UCSC Genome Browser track showing alignments of Illumina DNA-Sequencing reads from PCR amplicons for ORF9d (top) and ORF10 (bottom). The lower tracks show ribosome footprinting data. The region highlighted in green represents the location of the reverse primer, while red segments indicate the genomic position of the bands obtained by RT-PCR. The orange band indicates the genomic start of ORF9d. Under each genome browser track is displayed an IGV track of NRCeq untrimmed reads aligned in splice unaware mode (see **Materials and Methods**) to the NRCeq adapter concatenated to each assembled transcript.

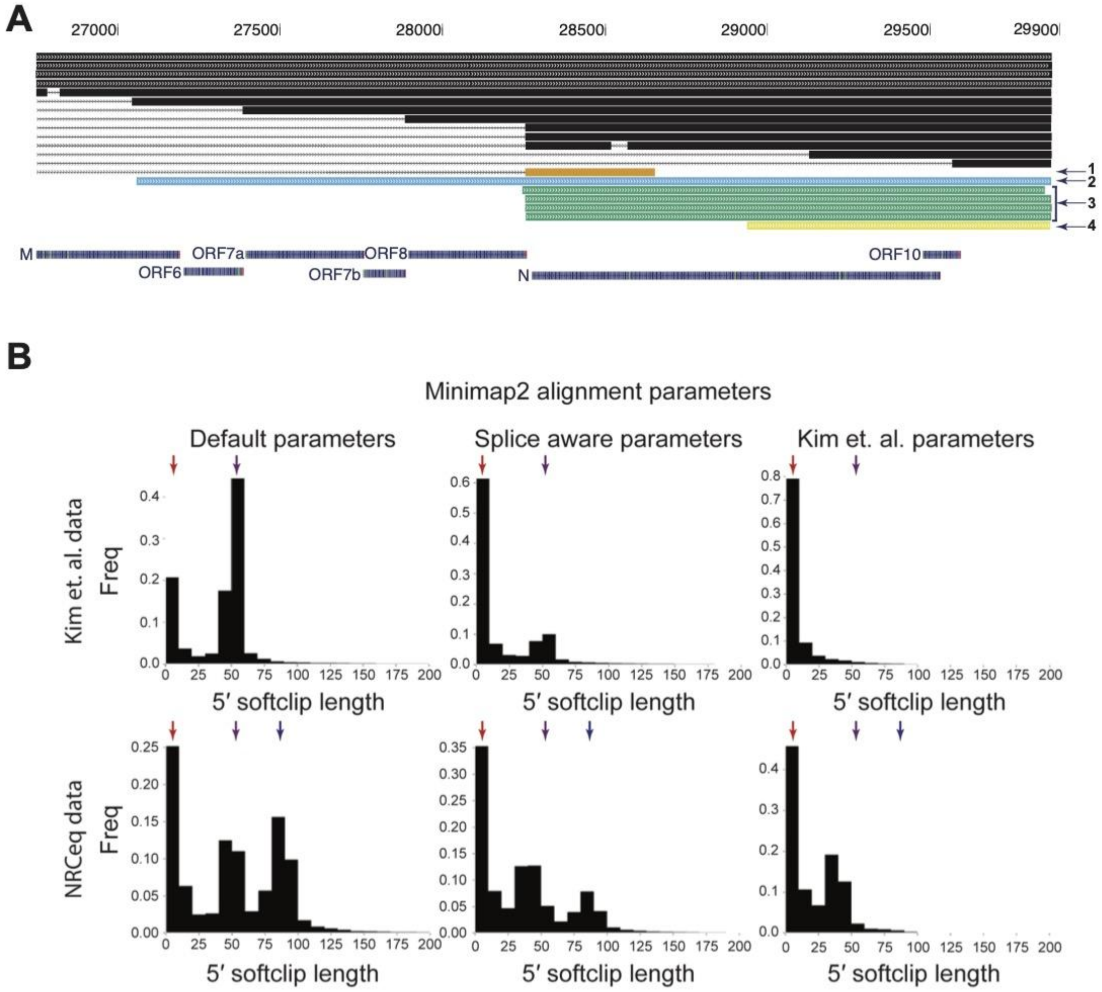


**Supplementary figure 6**. **Non-canonical transcript models.** (**A**) Non-canonical transcript model track from NRCeq assembly. Arrows indicate four types of non-canonical transcript models. (**B**) Frequency plots of lengths of softclipped sequences at the start site of all the genomic alignments of the datasets analyzed. The upper row distributions are calculated for Kim et al. dataset respectively with default long-reads parameters, splice-aware parameters and Kim et al. parameters. The bottom row represents the same analysis for NRCeq reads.

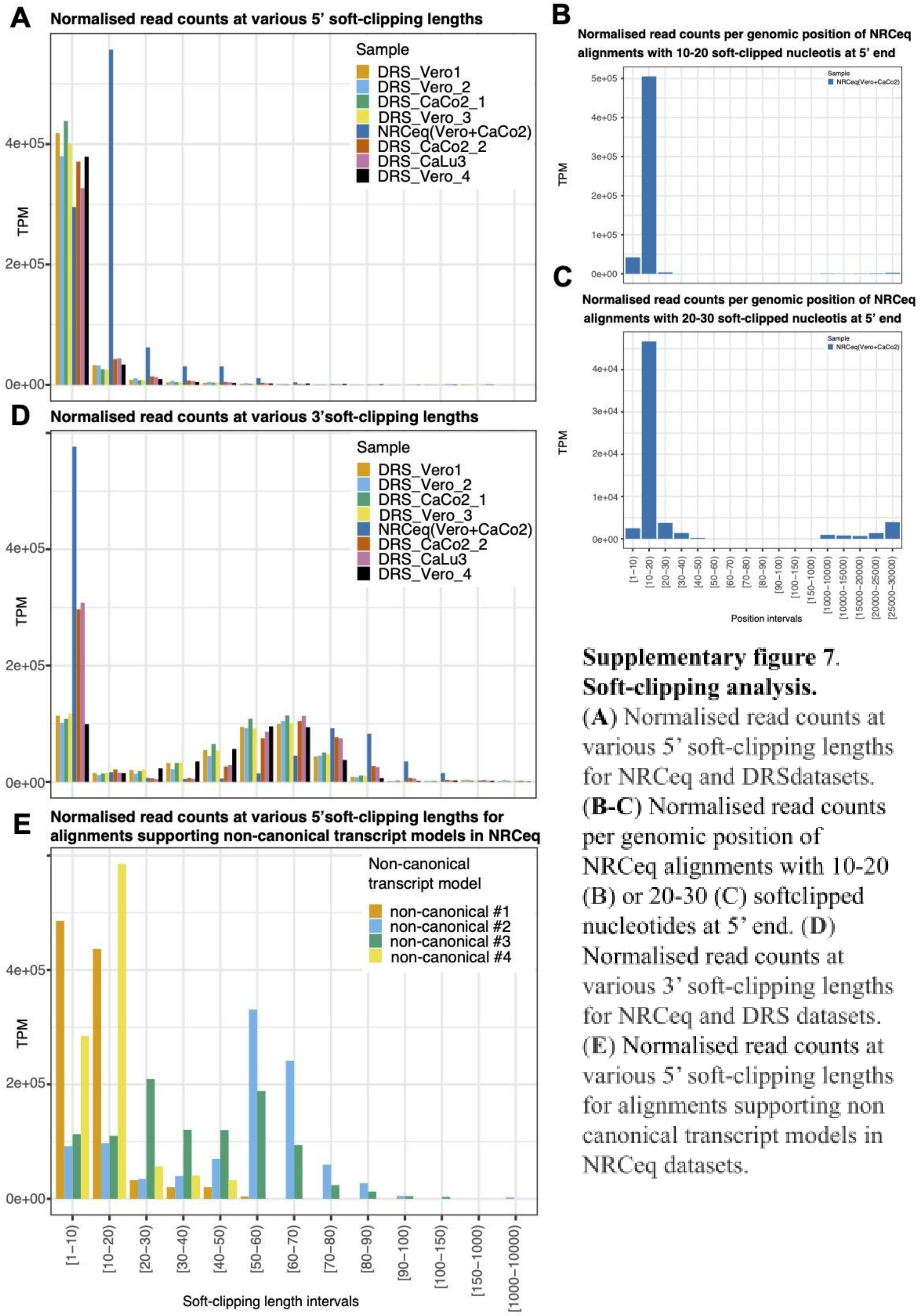


**Supplementary figure 7. Soft-clipping analysis.** (**A**) Normalised read counts at various 5' soft-clipping lengths for NRCeq and DRSdatasets. (**B-C**) Normalised read counts per genomic position of NRCeq alignments with 10-20 (B) or 20-30 (C) softclipped nucleotides at 5' end. (**D**) Normalised read counts at various 3' soft-clipping lengths for NRCeq and DRS datasets. (**E**) Normalised read counts at various 5' soft-clipping lengths for alignments supporting non canonical transcript models in NRCeq datasets.

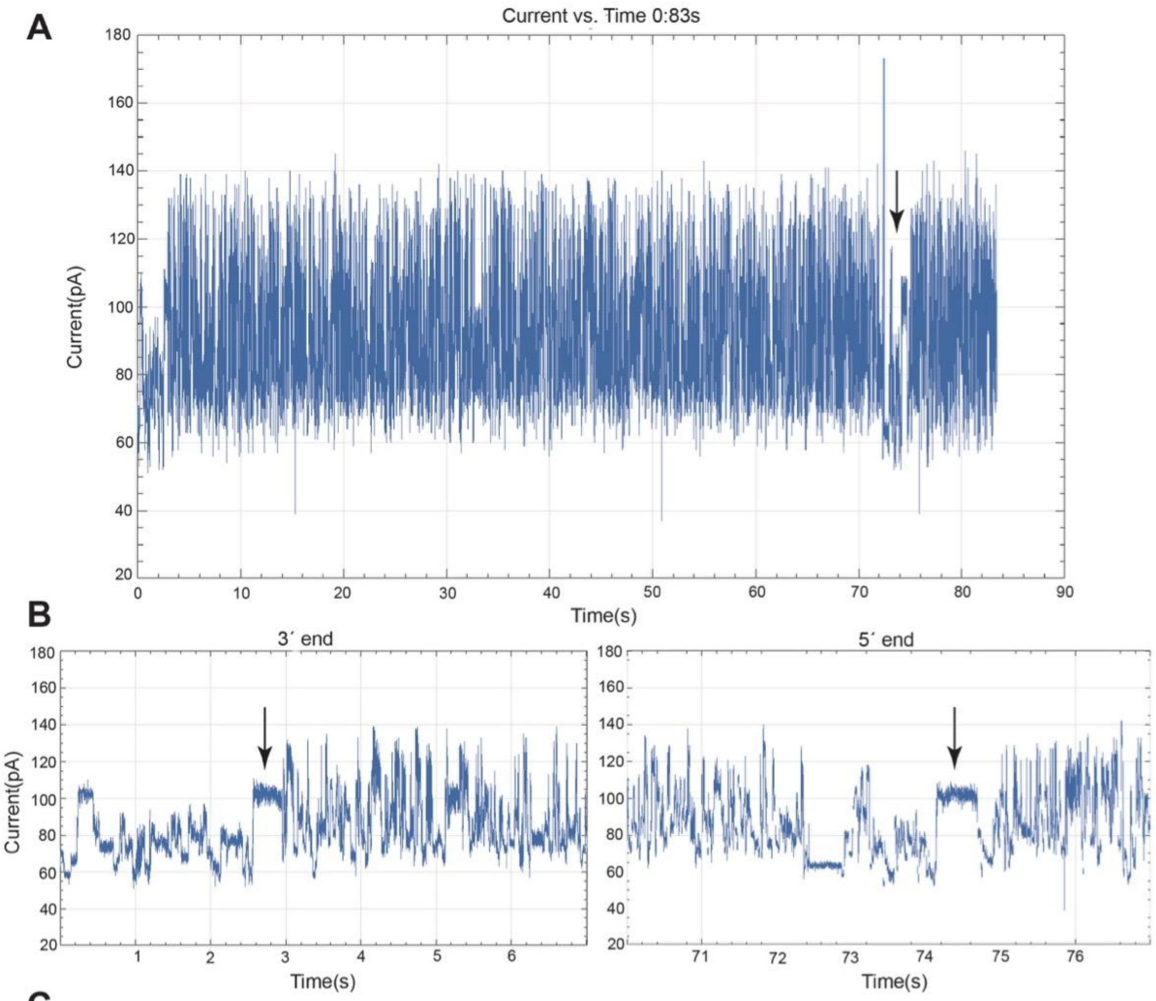


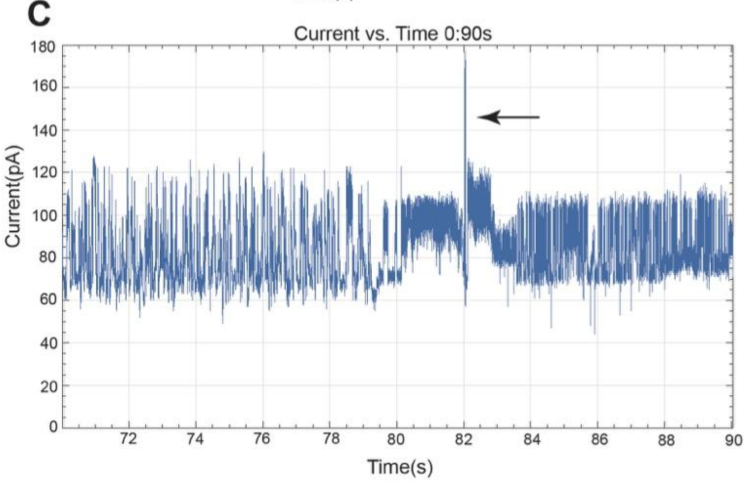


**BLAT to SARS-CoV-2 Reference**

| start | end | span of read | read length | span of reference |
|---|---|---|---|---|
| 289 | 6978 | 6689 | 7055 | 6811 |
| 14 | 3209 | 3195 | 7055 | 14 |
| 12 | 6762 | 6750 | 7055 | 12 |

**BLAT to HG38**

| start | end | span of read | read length | span of reference |
|---|---|---|---|---|
| 86 | 122 | 36 | 7055 | 34 |
| 105 | 134 | 29 | 7055 | 30 |
| 51 | 124 | 73 | 7055 | 38 |
| 107 | 135 | 28 | 7055 | 29 |
| 107 | 134 | 27 | 7055 | 28 |
| 41 | 69 | 28 | 7055 | 30 |
| 154 | 180 | 26 | 7055 | 27 |
| 76 | 100 | 24 | 7055 | 25 |
| 9 | 33 | 24 | 7055 | 25 |
| 78 | 104 | 26 | 7055 | 26 |
| 82 | 110 | 28 | 7055 | 28 |
| 100 | 123 | 23 | 7055 | 47 |
| 107 | 129 | 22 | 7055 | 23 |
| 100 | 122 | 22 | 7055 | 23 |
| 78 | 99 | 21 | 7055 | 22 |
| 43 | 64 | 21 | 7055 | 22 |
| 43 | 64 | 21 | 7055 | 22 |
| 78 | 99 | 21 | 7055 | 22 |
| 44 | 64 | 20 | 7055 | 21 |
| 107 | 127 | 20 | 7055 | 21 |
| 107 | 127 | 20 | 7055 | 21 |
| 137 | 156 | 19 | 7055 | 20 |
| 107 | 126 | 19 | 7055 | 20 |
| 45 | 64 | 19 | 7055 | 20 |

**Supplementary figure 8**. **Examples of signal artifices.** (**A**) Example of reads concatenation. The arrow indicates the poly(A) tail of the second read linked to the 5’ of the first read. (**B**) Close view of the 5’ and 3’ end of the read represented in ( **A**). Arrows indicate the poly(A) tails of concatenated reads. (**C**) On the left, example of a read getting stuck in the pore and “kick out” attempt by current polarity inversion. On the right, BLAT of the softclipped sequence of the read to the HG38 and SARS-CoV-2 genome.

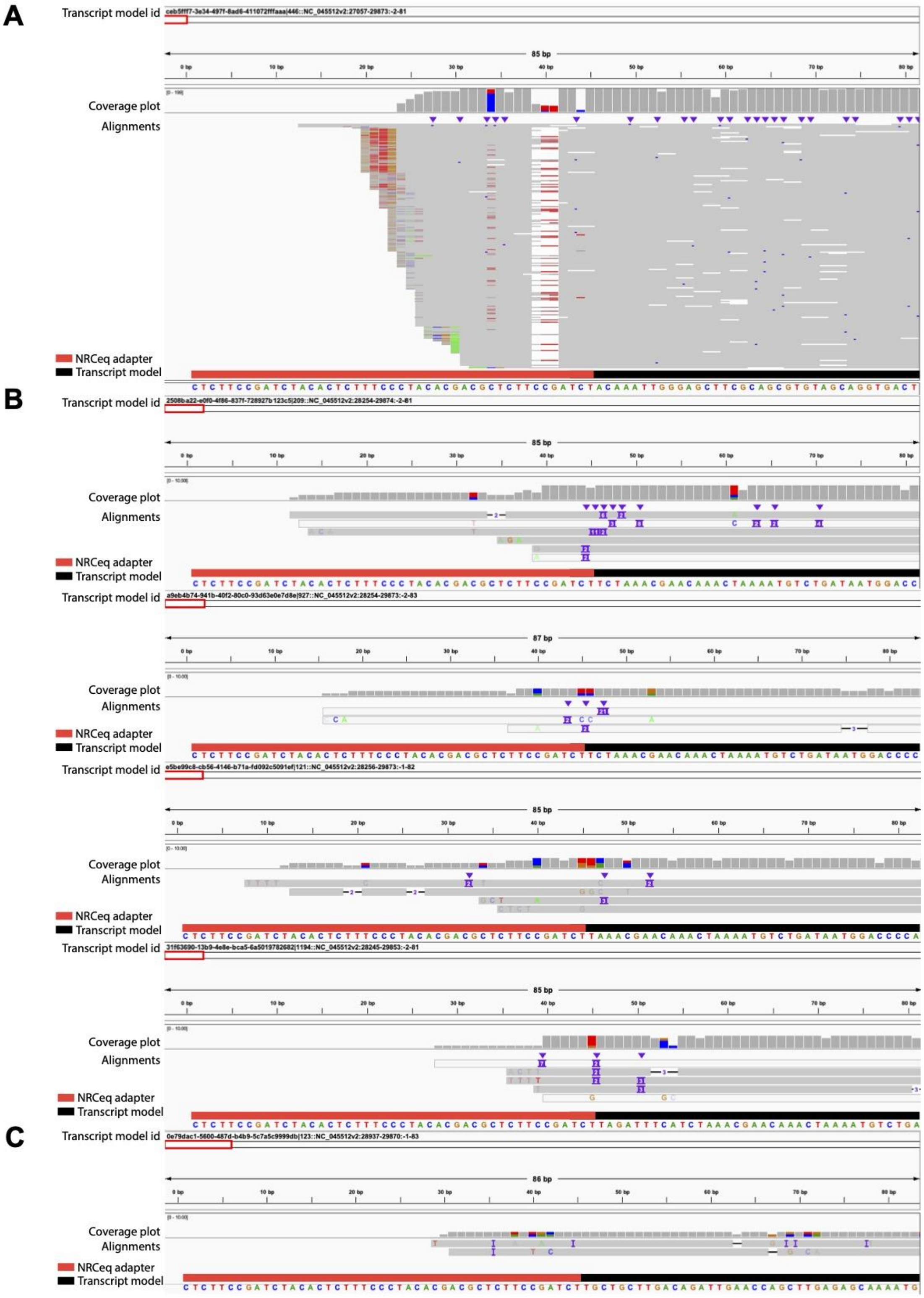


**Supplementary figure 9**. **Non-canonical transcript models alignments.** Tracks from IGV[42]. (**A**) NRCeq untrimmed reads aligned to the non-canonical transcript model sequence #2 preceded by the adapter sequence. The gap at 40bp represents the miscalling due to the PEG-spacer. (**B**) NRCeq untrimmed reads respectively aligned to the four non-canonical transcript model sequences belonging to group #3 preceded by the adapter sequence. The gap at 40bp represents the miscalling due to the PEG-spacer. (**C**) NRCeq untrimmed reads aligned to the non-canonical transcript model sequence #4 preceded by the adapter sequence. The gap at 40bp represents the miscalling due to the PEG-spacer.

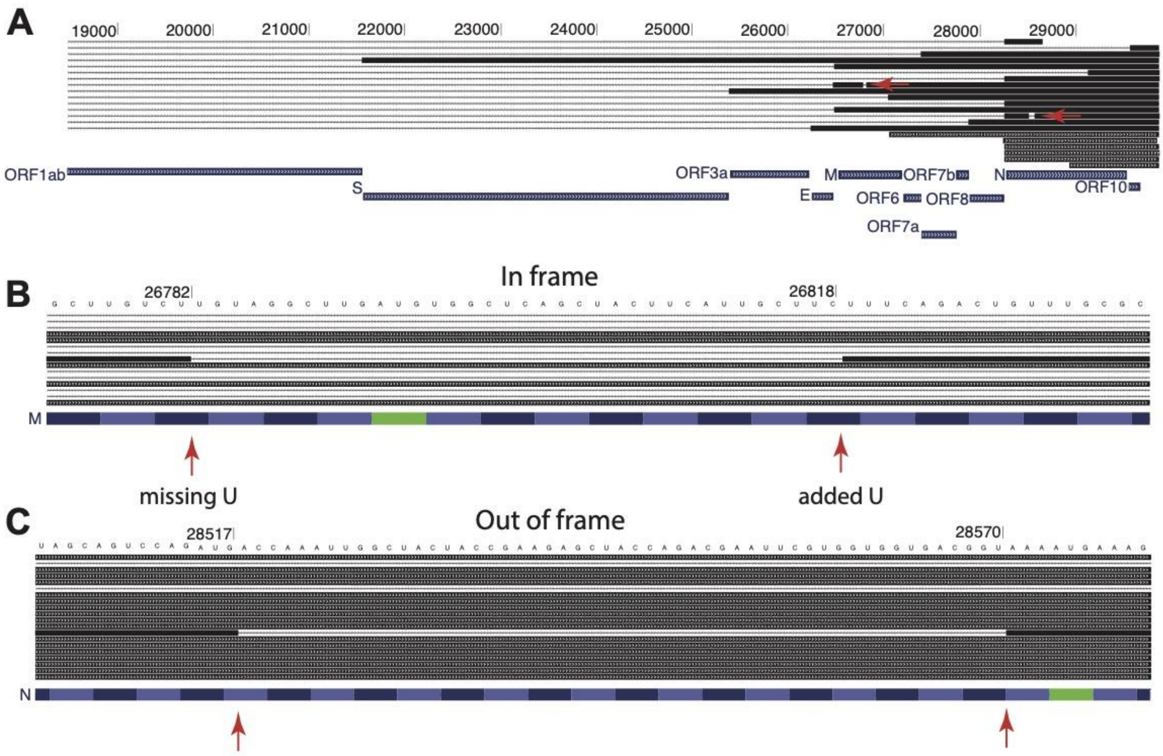


**Supplementary figure 10**. **Deletion analysis.** Tracks from the UCSC Genome Browser. (**A**) Overview of the region of the assembly where the deletions are localized. Red arrows point to two deletions. (**B-C**) Close view of the deletions in (**A**). Red arrows indicate the start and the end of the deletions. (**B**) is in-frame, a uridine has been mapped on the end of the deletions but is missing at its start. (**C**) is out of frame.

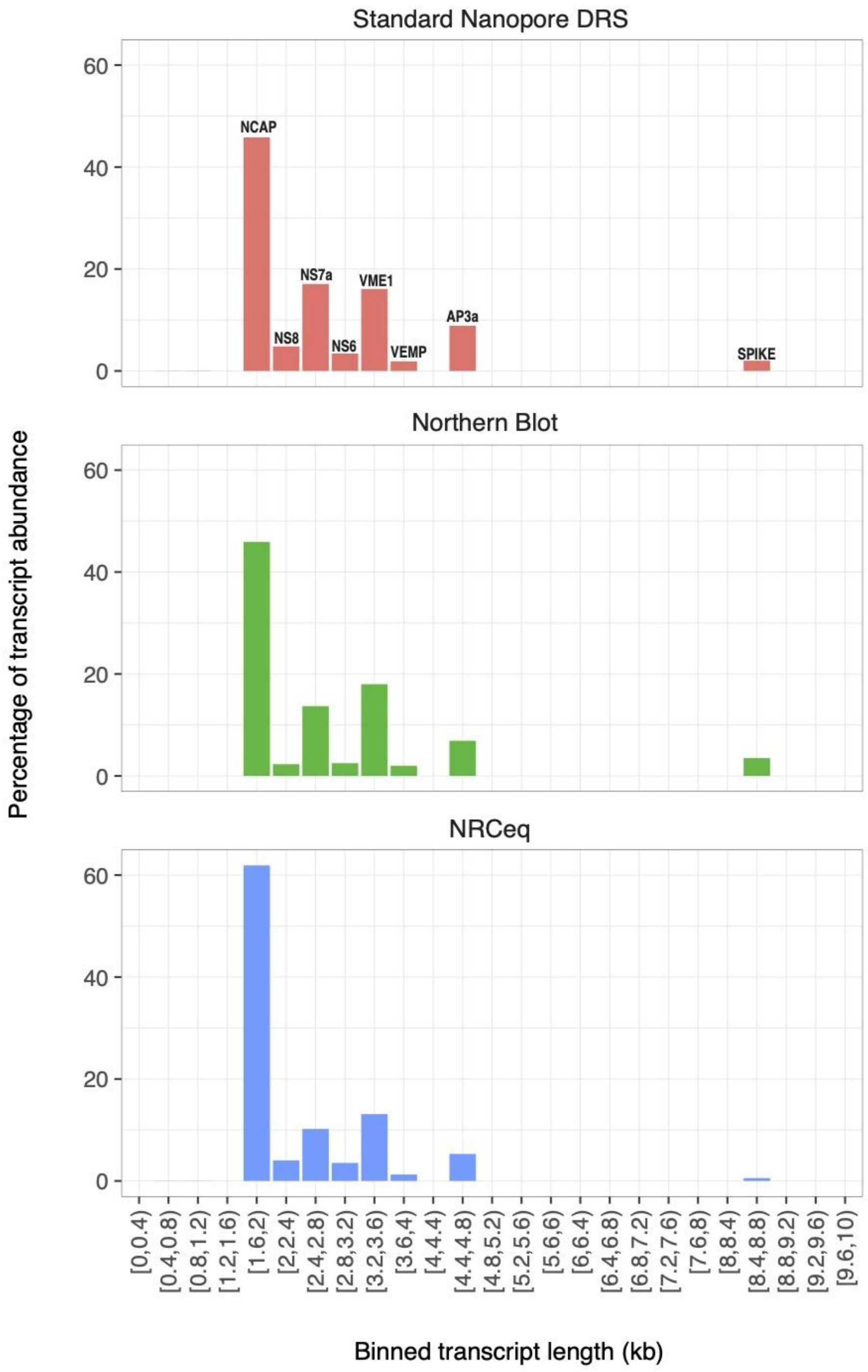


**Supplementary figure 11**. **Comparison of ORF quantification.** Quantification of the ORFs performed by Standard direct RNA Nanopore Sequencing, Northern Blot[22] and NRCeq.

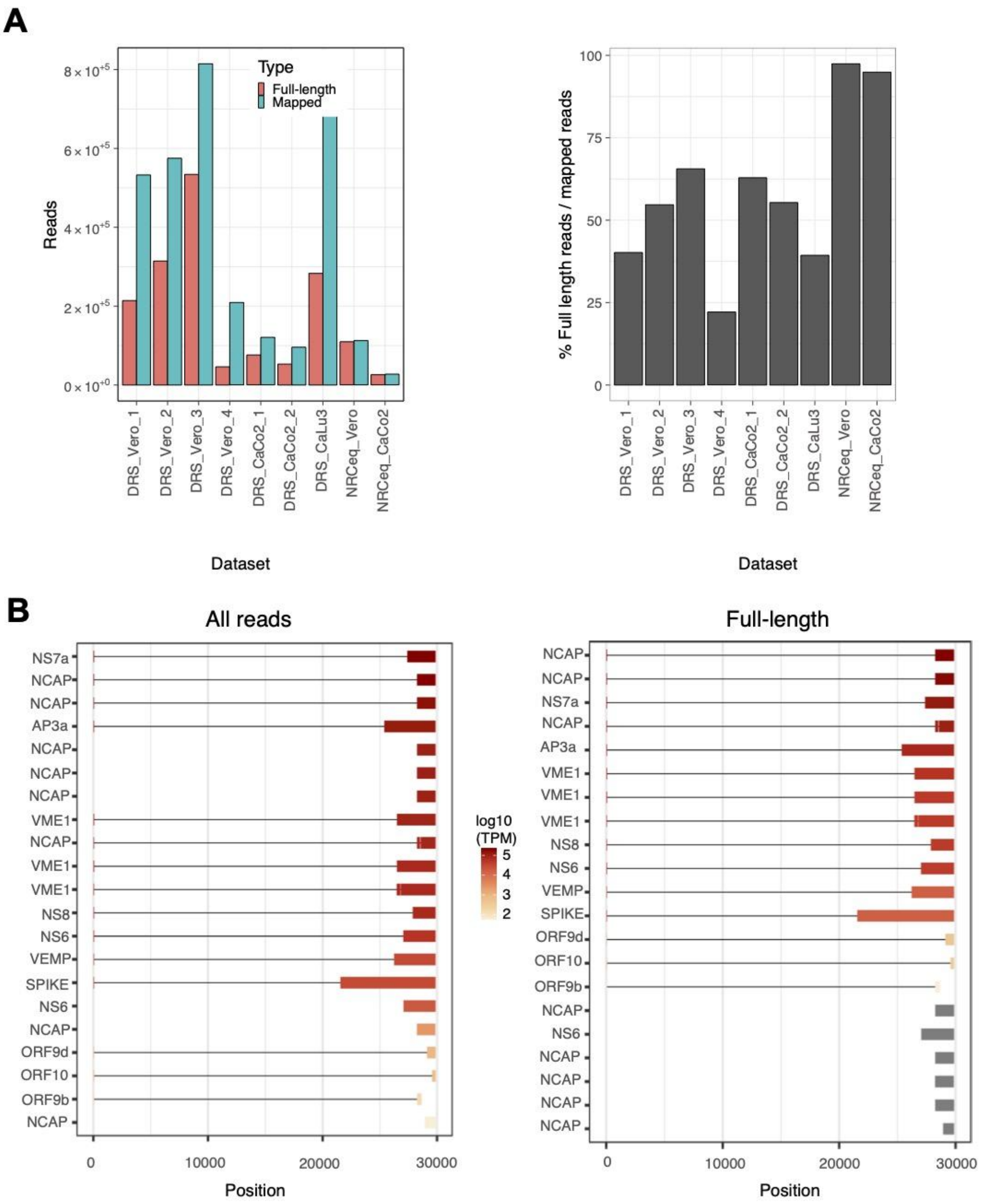


**Supplementary figure 12**. **Full-length approach allows to filter out a large amount of incompletely sequenced molecules**. (**A**) Number (left) and percentage (right) of reads mapped to the SARS-CoV-2 genome (Mapped) and filtered according to the full-length approach (Full-length) for each sample. (**B**) Structure and expression of the transcript models in the assembly. On the left, the analysis includes all reads of each sample; on the right, reads have been filtered according to the full-length approach (Panel on the right is the same as **Fig. 3C**).

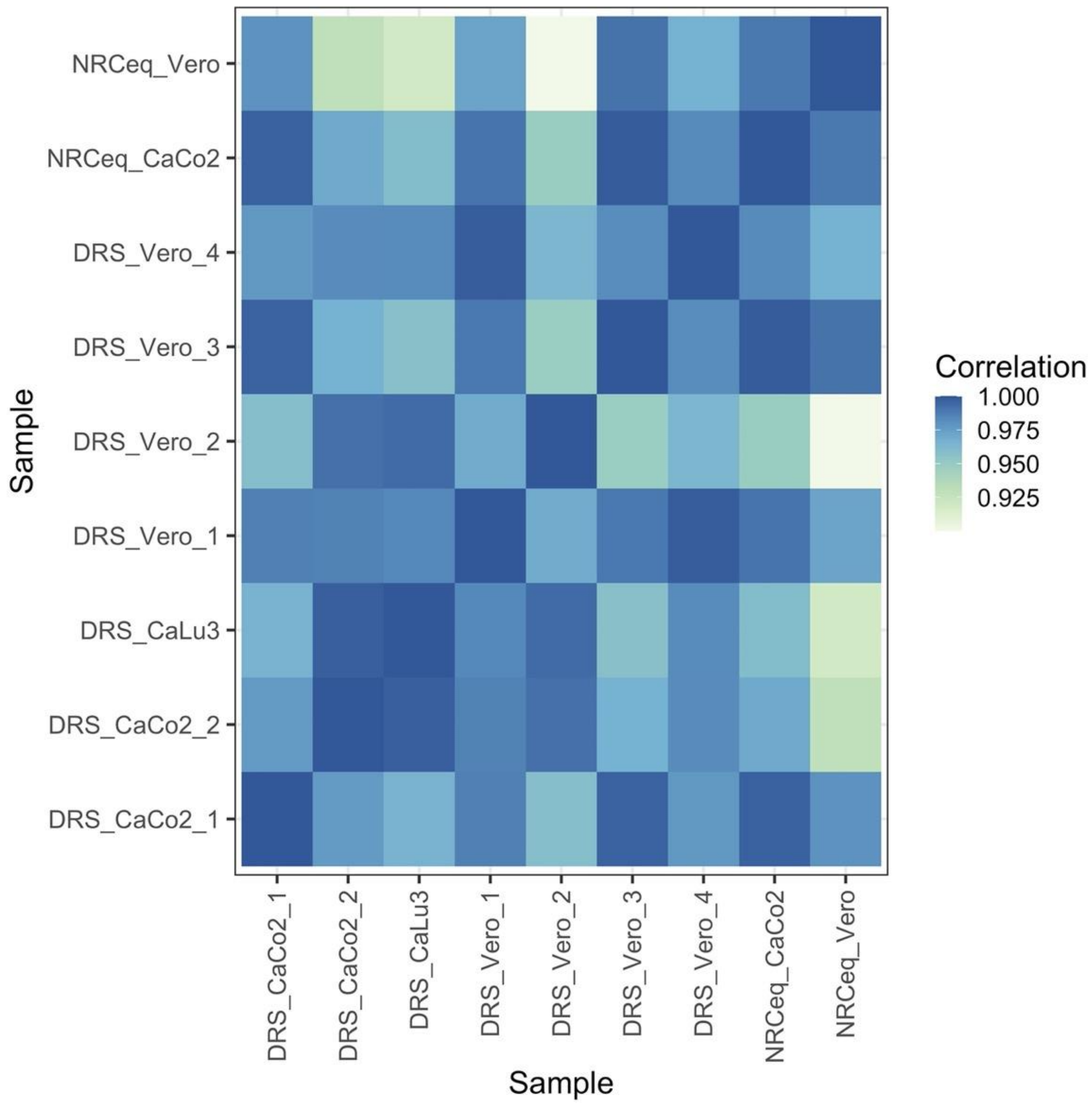


**Supplementary figure 13**. **Sample correlation matrix**. Sample expression Pearson correlation matrix. Tpms have been grouped per ORF.

### Supplementary Tables

**Supplementary Table 1:** Datasets generated as part of this study
**Supplementary Table 2**: Annotation of NRCeq Start Sites peaks in BED format
**Supplementary Table 3**: Annotation (BED format) and sequence (FASTA format) of the NRCeq transcript models.
**Supplementary Table 4**: Ranking of SARS-CoV-2 ORFs according to their expression levels obtained with DRS, NRCeq and Northern blot.

#### Supplementary Track

A custom UCSC Genome Browser hub with data generated in this study is available at: https://doi.org/10.6084/m9.figshare.17061377
The hub includes the following tracks:

- NRCeq assembly
- NRCeq alignments 5' peaks (as in **Supplementary Figure 3)**
- Total coverage of NRCeq datasets (plus strand)
- Coverage of the first nucleotide of each alignment of NRCeq datasets (plus strand)
- Total coverage of standard DRS datasets (plus strand)
- Coverage of the first nucleotide of each alignment of standard DRS datasets (plus strand)
- Coverage of Illumina reads (from the short sgRNA amplicons) supporting a junction with the TRS-L

### Supplementary Information

**The presence of negative strand alignments in SARS-CoV-2 datasets**
The coverage for NRCeq and standard DRS datasets is shown only for the positive strand alignments, as reads mapping to the negative strand are found in a very small quantity (**Suppl. Fig. 2**). In fact, DRS sequences mRNAs from their poly(A) tail and the transcription of SARS-CoV-2 proceeds using negative intermediates to produce positive mature sgRNAs. These features lead to a much lower amount of negative strand alignments in the datasets analysed which can subsequently be overlooked.

**The amount of soft-clipping at the 5' and 3' ends is influenced by minimap2 parameters**
Current long-read aligners are mainly developed for mammalian sequences and the combination of parameters used can deeply influence the outcome of the analysis. In particular **Suppl. Fig. 6B** shows that the distribution of 5' softclip length depends on minimap2[18] parameters. We tested three combinations of parameters: default DRS minimap2 conditions, "splice aware" conditions (--splice) and parameters reported in Kim et al.[4] (see **Materials and Methods**). We computed the frequency and the amount of softclipping at the

5’ of the alignments for each condition, both for data from Kim et al. and our own NRCeq data.
For the Kim et al. dataset, the default and “splice aware” parameter combinations recovered two peaks: the peak of softclipped length between 0 and 10 corresponds to 0-10 terminal nucleotides of the molecule that are often miscalled in DRS; the other peak originates from soft-clipped 5’ UTR sequences. These sequences can be soft-clipped by the aligner for different, often combined, reasons. TRS-L are short sequences which share similarities with TRS-B sequence, something which - if combined with an incomplete sequencing or high number of mismatches in the whole 5’ UTR sequence - can lead the aligner to choose to softclip them instead of opening a gap after the TRS-B because of its entropic cost. Indeed, this second peak almost disappears for Kim et al. conditions, as they fine tune both the cost of gap opening and of its extension.
For NRCeq untrimmed datasets instead, with the default and “splice aware” parameter combinations, the recovered peaks were three. The first (0-10), due to the miscall of the terminal nucleotides , the second (25-50) due to the softclipping of the NRCeq adapter and the third (75-100) due to the softclipping of the combination of adapter and 5’ UTR with low mapping score.
In both the Kim et al. and NRCeq datasets, the peaks deriving from the softclipping of the 5’ UTR are prevalently removed using the parameters proposed by Kim et al., which lower the penalty score of opening and extending a gap.

**Non-canonical sgRNAs analysis**
Distinguishing between real non-canonical sgRNAs and artefacts requires a deep analysis on the alignments that support each transcript model. Artefacts could be generated by inaccuracies in the basecalling, trimming and especially mapping. Indeed, when we utilize an aligner that has been tuned principally for mammalian sequences, such as minimap2[18], the combination of mapping parameters can deeply influence the presence of artefacts (**Suppl. Fig. 6B**), especially in the case of the SARS-CoV-2 transcriptome which is composed of nested transcripts.
This is why alignments supporting non-canonical transcript models have been manually inspected (**Suppl. Fig. 6A**) to assess if they were real molecules or artefacts deriving from incorrect mapping or assembling.
Our analysis shows that:

- non-canonical transcript model #1 is truncated at the 3’ in correspondence of a stretch of As, which can be interpreted by the sequencer as a polyA tail and consequently basecalled as a truncated molecule;
- non-canonical transcript models #4 is supported by alignments which are canonical but that are displayed as non-canonical due to the softclipping of the TRS-L. Indeed these reads have different leader-body fusion sites due to the scoring method of minimap2. When the region at the 3’ is short and the ORF sequence has similarities with the 5’ UTR, the aligner is not able to recover the exact TRS-L/B junction site and gives different results according to the mismatches that it encounters. This is why this transcript model includes reads supporting ORF9, which are not sufficient to form a transcript model themselves due to Pinfish parameters;
- non canonical transcript models #2 and group #3 are supported by alignments that include possible TRS-L softclipping events and potential real non-canonical sgRNAs.

To distinguish between artefacts and real non-canonical sgRNAs we have established two criteria. A read is defined as a real non-canonical sgRNA if:

- the first 60 nucleotides of the read do not map to the 5’ UTR but map with at least 70.0% identity to a downstream sequence;

- the 5’ of the read overlaps with one of the 5’ m7G cap sites identified by NRCeq (**Suppl. Fig. 2**).

Applying this criteria we found ~50 reads supporting these transcript models, providing support for the existence of genuine capped non canonical sgRNAs.
This puts in evidence the biases that the previous works with standard dRNA Nanopore Sequencing were suffering from, as they identified a large amount of non-canonical leader-to-body junctions that in many cases could be hardly distinguished from incomplete sequencing or degradation fragments molecules.

**Soft-clipping analysis reveals additional information on the transcript origin**

We have performed a softclipping analysis on the 3’ and 5’ of alignments of standard DRS and NRCeq datasets.

For the soft-clipping at the 3’ end of the alignments, we have observed that the number of softclipped bases was between 10 and 20 nucleotides for most of the reads from all of the datasets. There was a supplementary peak between 50 and 80 bases softclipped (**Suppl. Fig. 7D**).

The soft-clipping at the 5’ end has already been described in the section **Results**, but further information on the alignments origin can be obtained by manual inspection of the soft-clipped sequences.
First of all, we noticed a very low percentage of alignments with long softclipping (1kbp) which we investigated by hand to understand their origin in order to understand if there could be ligation events between human and viral molecules.
We can divide our observations in two case studies: the first is an event in which we observe an artificial concatenation of two viral molecules. This can be originated from two molecules entering one after the other in the pore which are spatially too close to be identified as distinct entities by the basecaller (**Suppl. Fig 8A-B**). In this case the second viral molecule is entirely softclipped.
The other case occurs when we observe that part of the softclipped sequence maps to the viral genome and part to the human genome. This artefact can originate when a viral molecule gets stuck in the pore, which inverts the polarity of the current in order to kick it out causing a miscalling in the downstream signal, that, by chance, makes the sequence alignable to the human genome. This can be also seen by the aligning BLAT scores in **Suppl. Fig. 8C**.
Another observation that has been made about softclipping is the possibility of creating artefacts in the form of non-canonical sgRNAs. As previously described, the region of the 5’ is quite short and eventually rich in mismatches, due to miscalling of the terminal 10 nucleotides or to possible modifications. This makes this region more prone to being softclipped, which in some cases might be more entropically convenient than opening and extending a gap for 2kbp. This issue can generate artefacts in the mapping which can be displayed as non-canonical alignments. Indeed when analyzing the transcript abundance at various 5p softclip length for alignments supporting non-canonical transcript models in NRCEq datasets we observe a peak between 50 and 80 nucleotides for non-canonical transcript models #2 and #3. These reads were also mapped to the viral genome through local alignment tools ( algorithm with parasail, see **Materials and Methods** - data not shown) and confirms our hypothesis of a 5’ softclipped 5’ UTR for most of them. Beside this, still ~50 reads that support non-canonical transcript models did not show a TRS-L or an untrimmed adapter in the softclipped sequence, and this is why we performed a specific analysis for non-canonical transcript models (see **Non-canonical sgRNAs**).

**Full-length approach**

To avoid possible quantification errors due to RNA degradation or incomplete sequencing, we filtered the reads of every dataset that we wanted to quantify against our assembly using a full-length approach. This criteria defines full length reads as those having the 5’ in an interval of 100 nucleotides from on the 5’ of each transcript of the NRCeq assembly. The vast majority of the NRCeq reads are already full-length (**Suppl. Fig. 12**) while this approach is mostly useful when dealing with standard DRS datasets as it helps to give a better quantification of each isoform. It is clear how non-canonical transcripts recovered by the assembly have a much lower expression compared to the canonical ones (**Suppl. Fig. 12C**)
